# Connectomics of the *Octopus vulgaris* vertical lobe provides insight into conserved and novel principles of a memory acquisition network

**DOI:** 10.1101/2022.10.03.510303

**Authors:** Flavie Bidel, Yaron Meirovitch, Richard Lee Schalek, Xiaotang Lu, Elisa Catherine Pavarino, Fuming Yang, Adi Peleg, Yuelong Wu, Tal Shomrat, Daniel Raimund Berger, Adi Shaked, Jeff William Lichtman, Binyamin Hochner

## Abstract

We present the first analysis of the connectome of the vertical lobe (VL) of *Octopus vulgaris*, a brain structure mediating acquisition of long-term memory in this behaviorally advanced mollusk. Serial section electron microscopy revealed new types of interneurons, cellular components of extensive modulatory systems and multiple synaptic motifs. The sensory input to the VL is conveyed via ~1,800,000 axons that sparsely innervate two parallel and interconnected feedforward networks formed by the two types of amacrine interneurons (AM), simple AMs (SAMs) and complex AMs (CAMs). SAMs make up 89.3% of the ~25,000,000 VL cells, each receiving a synaptic input from only a single input neuron on its non-bifurcating primary neurite, suggesting that each input neuron is represented in only ~12 SAMs. This synaptic site is likely a “memory site” as it is endowed with LTP. The CAMs, a newly described AM type, comprise 1.6% of the VL cells. Their bifurcating neurites integrate multiple inputs from the input axons and SAMs. While the SAM network appears to feedforward sparse “memorizable” sensory representations into the VL output layer, the CAMs appear to monitor global activity and feedforward a balancing inhibition for “sharpening” the stimulus-specific VL output. While sharing morphological and wiring features with circuits supporting associative learning in other animals, the VL has evolved a unique circuit that enables associative learning based strictly on feedforward information flow.

## INTRODUCTION

The advanced learning capabilities of cephalopod mollusks like *Octopus vulgaris* are comparable to those in some vertebrates (Sanders 1975; Mather 1995; Zarrella et al. 2015; Schnell et al. 2016; Hanlon and Messenger 2018). As the last common ancestor of cephalopods and vertebrates existed more than 500 million years ago (Packard 1972; Vitti 2013), the octopus provides a unique opportunity to study the independent evolution of nervous systems subserving sophisticated cognitive abilities (Hochner et al. 2006; Shomrat et al. 2015; Hochner and Glanzman 2016; Turchetti-Maia et al. 2017). The vertical lobe (VL), the highest structure in the hierarchy in the central nervous system of the octopus, is regarded as a high order integrative center involved in visually evoked associative learning and memory acquisition (Boycott and Young 1955; Sanders 1975; Young 1979; Fiorito and Chichery 1995; Hochner 2010, 2012, 2013; Shomrat et al. 2015; Shigeno et al. 2018; Turchetti-Maia et al. 2017). It is exceptionally large, containing 25 million neurons, about half the neurons in the central nervous system. This number is not much smaller than the number of neurons in the human hippocampus (~40 million, West and Gundersen, 1990) and about 4 orders of magnitude greater than the number of neurons in the mushroom body (MB) of *Drosophila* (e.g. Aso et al., 2014) and three orders of magnitude greater than that in the locust MB (Jortner et al. 2007).

Extensive neuroanatomical data on the VL were provided by the work of J. Z. Young and E. G. Gray in the 1960-70’s using light and electron microscopy (Gray and Young 1964; Gray 1970; Young 1971). Based on estimated neuron numbers they proposed that the VL network is extremely simple, composed of only two types of morphologically distinguishable neurons: 25 million small amacrine interneurons (AM) and 65,000 large efferent neurons (LN). The main sensory input to the VL is conveyed via 1.8 million axons projecting from the median superior frontal lobe (SFL). These axons compose the SFL-VL tract that runs in the outer neuropil of the five VL lobuli. Each of these axons makes *en passant* synapses with a then unknown number of the AMs. These 25 million AMs converge sharply at a ratio of about 400:1 to innervate the LNs, whose axons were assumed to be the only output of the VL. Young pointed out that the pattern of SFL innervation of the AMs resembles the connectivity matrix in other learning and memory networks like the insect MB and the mammalian hippocampus (Young 1995).

This gross connectivity scheme provided insights that guided more recent physiological studies on the VL. Physiological experiments in the brain-slice preparation (Hochner et al. 2003, 2006) revealed that the *en passant* SFL-to-AM innervation is glutamatergic. It shows a robust activity-dependent NMDA-independent long-term potentiation (LTP) of glutamate release from the SFL presynaptic terminals (Hochner et al. 2003). The AM-to-LN connections are excitatory cholinergic synapses (Shomrat et al. 2011). Recently it was discovered that LTP at the SFL-to-AM synapse is maintained by a novel nitric oxide (NO)-dependent ‘molecular switch’ mechanism in which NO switches ON NO-synthase (NOS) through a positive feedback loop of NO-dependent NOS activation (Turchetti-Maia et al. 2018).

Although Gray and Young provided a detailed description of the main neuronal elements constituting the VL network, the inability to reconstruct individual neurites limited the possibility of providing a detailed circuit connectivity. Indeed, their observations reported unclassified cell types, thus suggesting that the VL network is likely composed of more than the 3 neuronal elements they characterized (SFL, AMs and LN). Previous findings in our group also hinted that the VL is more complex. Firstly, intracellular recording from LNs revealed that some receive inhibitory inputs (Shomrat et al., 2011). Secondly, a recent wide-ranging histolabeling study suggested that the AMs may be divided into at least two groups, one cholinergic and the other GABAergic (Stern-Mentch et al. 2022).

To understand the octopus VL circuitry at a higher resolution the current study utilizes recent advances in our ability to elucidate synaptic wiring diagrams at the nanoscale level using electron microscopy (EM). Widely applied to various species and tissues this methodology has contributed much to understanding the organization of neural networks (Kasthuri et al. 2015; Lee et al. 2016; Morgan et al. 2016; Ryan et al. 2016; Motta et al. 2019; Li et al. 2020; Macrina et al. 2021; Meirovitch et al. 2021; MICrONS Consortium et al. 2021; Verasztó et al. 2020; Witvliet et al. 2021; Shapson-Coe et al. 2021). Here we sought to produce a synapse-level wiring diagram of the *Octopus vulgaris* VL from serial EM by tracing out all pre- and postsynaptic partners of dozens of main afferents of the VL (SFL projecting axons) and AMs. Indeed, although we found that the gross anatomical organization of octopus VL fits the previous anatomical and physiological model described above, the nanoscale analysis revealed new types of neurons and connections that contribute profoundly to a deeper understanding of the functional organization of the octopus VL and how it implements associative learning and memory.

Our data show that, firstly, in contrast to what was previously thought, the AM interneurons are segregated into two distinct groups. The majority (~89.3 %) of the VL cells bodies are simple AMs (SAM) that send a single non-bifurcating neurite into the neuropil. A small fraction (~1.6 %) forms a very small group of newly discovered complex AMs (CAM) with a branching dendritic tree. Secondly, we found that the SAMs receive only one synaptic input from a single SFL axon. We propose that this synaptic connection may serve as a “memory site” as it shows a robust activity-dependent LTP following tetanization of the SFL axonal tract (Shomrat et al 2011). Thirdly, the segregated SFL-to-SAM expansion format suggests that each of the 1.8 million SFL neurons is represented extremely sparsely in only around 12 of the 22.3 million SAMs. Fourthly, in sharp contrast to the SAMs, the CAMs integrate dozen to hundreds of inputs, suggesting that these interneurons integrate broader incoming SFL activity. As the SAMs and CAMs converge onto the same output layer of LNs processes, our findings suggest the VL appears to be comprised of two interconnected feedforward networks - one excitatory, originating from the SAMs (each representing one SFL) and a second, a parallel feedforward network, likely inhibitory, originating from the CAMs (each representing several SFLs). We demonstrated that distinct groups of SAMs and CAMs neurites fasciculate together to form a columnar organization in which each column constitutes canonical microcircuitry that may function as an “association module”, associating SFL sensory features. Lastly, we described an additional afferent to the VL and two extrinsic putative neuromodulation systems. These interact with the feedforward networks and thus are likely involved in regulating the activity state of the network and in conveying reward and punishments signals.

## RESULTS

### The VL dataset and analysis

The region of interest (ROI) is in the lateral lobule of the VL (**Figure 1A**). We cut, collected and imaged 892 serial sections (30nm section thickness) at 4nm/pixel to generate a traceable 3D image stack covering a volume of 260×390×27 μm (**Figure 1B; Figure 1-video supplement 1**; ‘connectome dataset’ available at https://lichtman.rc.fas.harvard.edu/octopus_connectomes).

**Figure 1.**
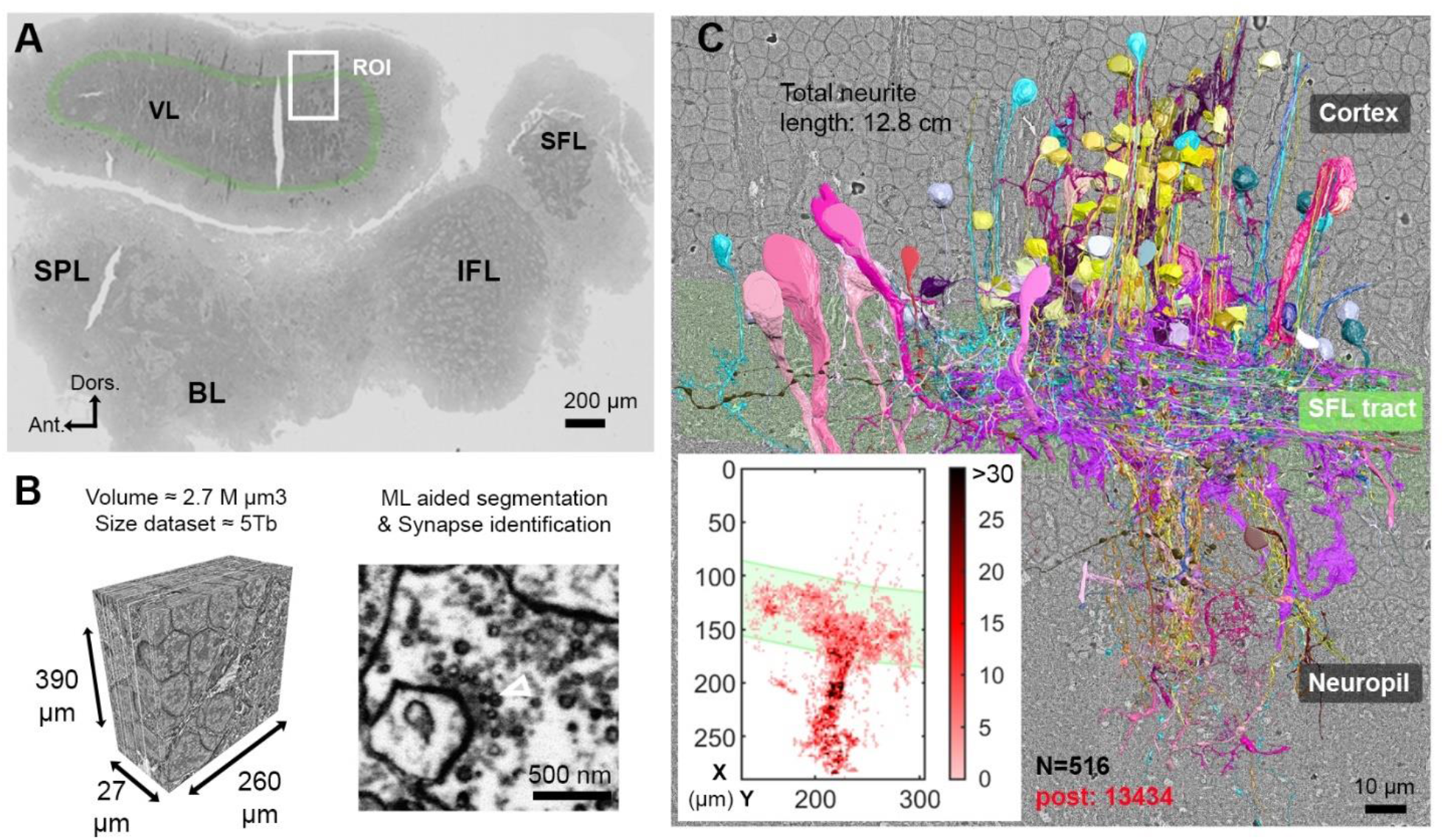
Reconstruction of octopus VL circuits using volume EM. (A) Low resolution image of the octopus central nervous system. The VL lies dorsally in the supraesophageal part of the central brain. The region of interest marked within it (ROI, white rectangle) was scanned with a scanning electron microscope (SEM) at 4 nm/px. (B) 30 nm sections were aligned in a traceable 3D stack (260×390×27μm. See also **Figure 1–video supplement 1**) for machine learning (ML)-aided by manual segmentation and synapse annotation. (C) The 516 reconstructed cell processes colored according to cell type (See **Figure 1-video supplement 2** for all cell types). Total neurite length of the reconstructed cells was 12.8 cm. Reconstructed cells are superimposed on a single EM image. Inset: spatial distribution of the 13434 postsynaptic sites annotated on the 516 cells presented as a heat-map. Abbreviations: BL, basal lobe; IFL, inferior frontal lobe; SFL, superior frontal lobe; SPL, subpedunculate lobe; VL, vertical lobe. See also **Figure 1–figure supplement 3** and **Figure 1–figure supplement 4**.

To classify the cells we used morphological features *(e.g.*, main neurite orientation, presence of varicosities, branching pattern), ultrastructural features *(e.g.*, vesicle size, mitochondria shape, cytoplasm staining, organelles) and connectivity assisted by the ‘blueprint’ established by Gray and Young (Gray and Young 1964; Gray 1970; Young 1971). Chemical synapses were identified based on features common to invertebrate synapses - the presence of a cluster of synaptic vesicles (sv) or ‘cloud’ associated with an active zone near the presynaptic membrane (Gray 1970; Ryan et al. 2016; White et al. 1986; Zheng et al. 2018; Meinertzhagen 2019; Witvliet et al. 2021). Synapses were annotated if this accumulation of sv persisted across at least three sections (> 90 nm). We classified synapses as fast chemical if the presynaptic component contained sv of 30-60 nm diameter and modulatory synapses when the synaptic component contained large vesicles (>60nm) (Messenger 1996; Leng and Ludwig 2008; van den Pol 2012; Witvliet et al. 2021). Consistent with previous studies (Gray and Young 1964; Gray 1970), we did not observe any obvious postsynaptic specialization *(i.e.*, postsynaptic density) at any chemical synapse and we found no evidence of gap junctions.

We reconstructed 516 cell processes with a total length of 12.8 cm. The reconstructed cell processes were classified into 10 cell types (**Figure 1–video supplement 2**) with 13,434 postsynaptic (**Figure 1C, heat map**) and 2,472 annotated presynaptic sites. Among these 10 cell types, only 7 were involved in the VL connectivity and therefore were analyzed in depth (see **Figure 1–figure supplement 3** for the 3 cell types not discussed here). For characterization of the 7 cell types, we reconstructed throughout the volume at least 30 processes per cell type and annotated all their pre- and postsynaptic sites. Additionally, a subset of cellular compartments and organelles *(i.e.*, cell bodies, nuclei and presynaptic varicosities) were separately segmented to allow morphological quantification. The core of our connectivity reconstruction included annotation and neurite reconstructions of all the synaptic partners of a subset of randomly selected ‘seed’ cells (Morgan et al., 2016): SFL axons (35 of 84 SFL axons reconstructed) and SAM cells (28 of 108 SAMs reconstructed).

We divided the analysis according to the three VL regions: 1) The outer cell body cortex; 2) the central neuropil, and 3) the SFL tract, or outer neuropil, where axons projecting from the SFL are arranged in a tract forming a ring around the outer zone of neuropil (Gray and Young, 1964; Young, 1971, Stern-Mentch et al 2022). Note that because most of the cells are not fully contained in the reconstructed volume, it is impossible to assess the full collection of synapses between pairs of neurons. We therefore used “synaptic site” to refer to a single anatomical release site or a single synapse.

### The VL connectome is characterized by two parallel and interconnected feedforward networks

**Figure 2** summarizes the 7 identified neuronal cell types and their connectivity (see also **Figure 2–figure supplement 1**). None of the 84 reconstructed SFL axons bifurcated within the ROI (~260 μm long) suggesting a very low rate of bifurcation of the SFL axons, if at all, as suggested by Young (1971). Each SFL axon innervated *en passant* two distinct sub-populations of AMs; a very large population of morphologically simple cells, which we termed ‘simple amacrine’ (SAM) and a small population of morphologically complex cells that we termed ‘complex amacrine’ cells (CAM) (**Figure 2 and Figure 3**).

**Figure 2.**
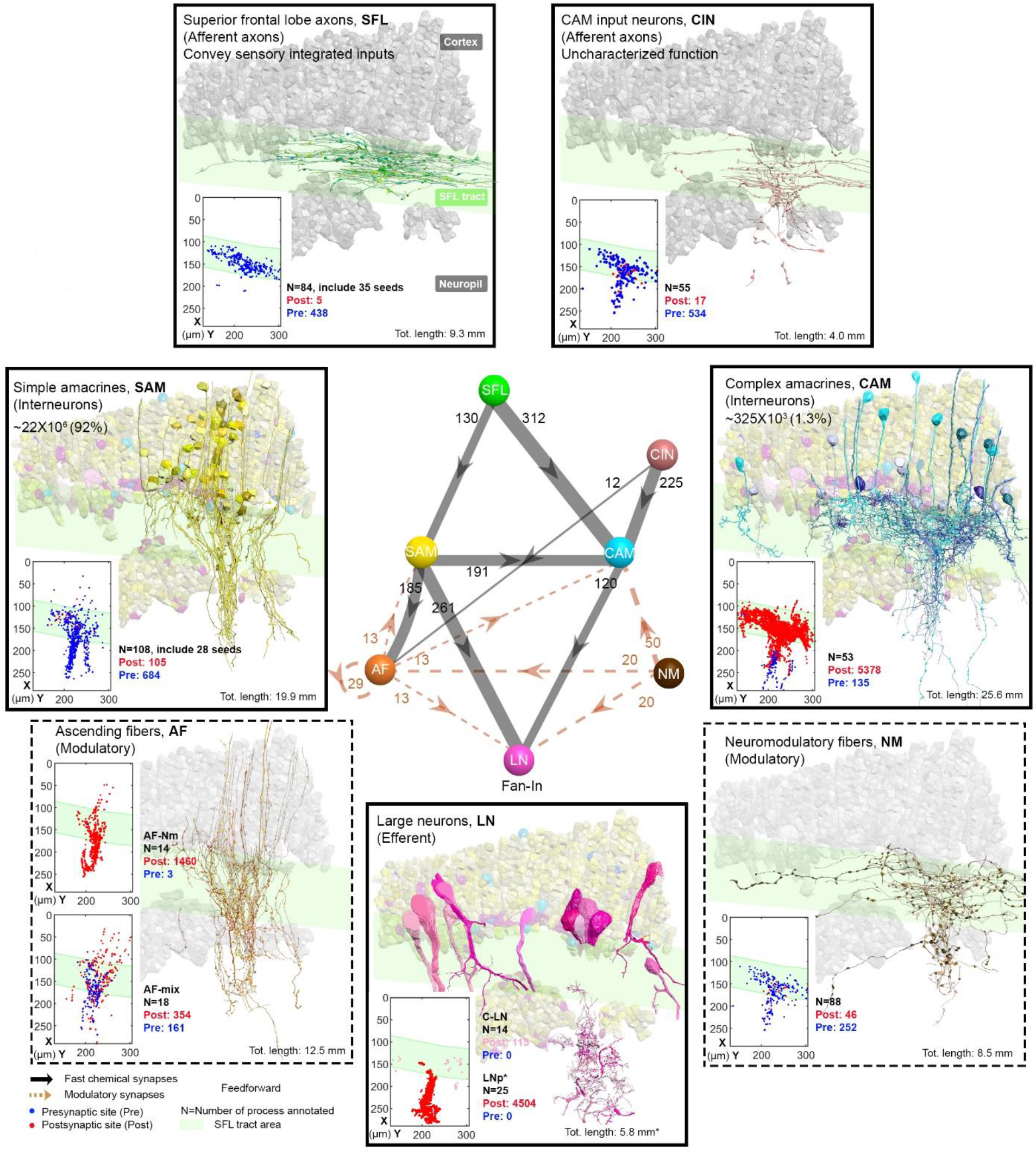
The 7 neuronal cell types classified in the ROI and their wiring diagram. The middle panel shows the wiring diagram, and the other panels depict the 7 neuronal cell types together with the spatial distribution of their postsynaptic sites (red puncta) and presynaptic sites (blue puncta). The approximate position of the SFL tract area (green) is shown for orientation and demarcating the outer from the inner neuropil. In the central panel each gray solid edge represents fast chemical synapses and the arrowhead the direction of these synapses; line width indicates relative number of synapses calculated from the number of connections annotated (noted next to each outgoing arrow). Orange dashed edges indicate chemical synapses observed with modulatory processes. Edges were represented only if more than 10 synapses were annotated. Note that the connectivity is strictly feedforward (p<0.05). See **Figure 2–figure supplement 1** for neurite level analysis.

**Figure 3.**
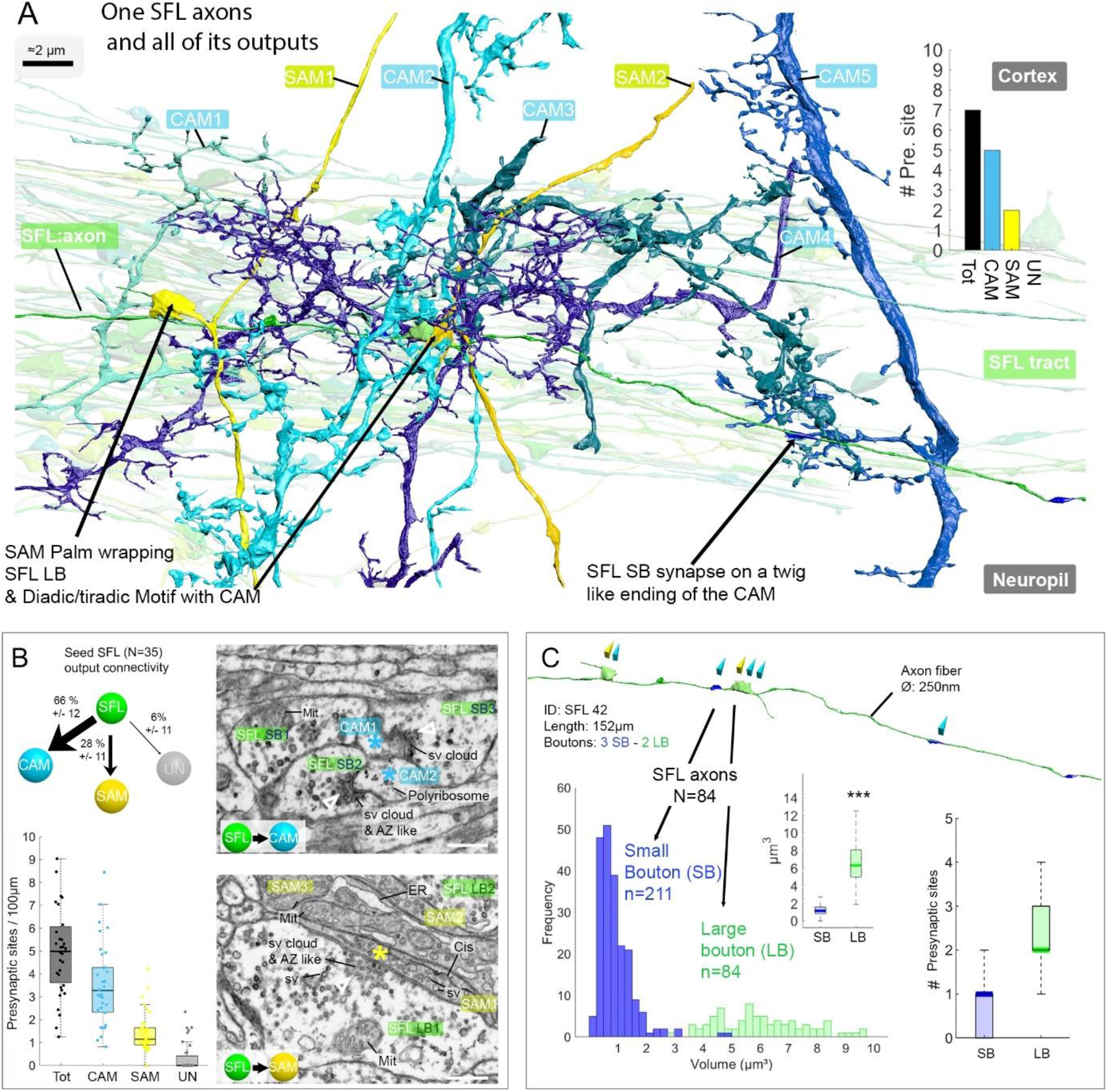
The SFL axons synapse onto the two populations of amacrine interneurons (SAMs and CAMs) via two distinct populations of presynaptic boutons. (A) Reconstruction of a single SFL axon (green) and its 7 *en passant* outputs within the reconstructed volume: 5 CAMs (various blue hues) and 2 SAMs (yellow). (B) Quantification of the synaptic outputs of the seed SFL axons (n=35) to SAMs (via large boutons; LB), CAMs (via small boutons; SB) and onto unidentified cell types (UN). Upper left shows distribution of connections in percent. On the right high resolution EM images of the SFL synaptic outputs onto SAM and CAM. 1nm/px, dwell time: 200 μs. Scale bar, 500 nm. Empty triangle and asterisk are respectively pre- and postsynaptic profiles. The whisker plot shows the relative density of the different presynaptic boutons along the 35 seed SFLs axons (C) Example of the distribution along an SFL axon segment of presynaptic outputs onto two SAMs (yellow triangles) and CAMs (light blue triangles). The histogram displays the bimodal distribution of bouton volumes of 84 SFL axons into two populations of small boutons (SB-blue) and large boutons (LB-green). The plot of presynaptic sites per SB and LB reveals their monosynaptic and multisynaptic nature, respectively. Asterisks indicate statistical significance *** p<0.001, **<0.01. Abbreviations: sv, synaptic vesicles; AZ-like, active zone-like; Mit, mitochondria, Cis, cisternae; tER, transport endoplasmic reticulum. See also **Figure 3–figure supplement 1** and **Figure 3–video supplement 2**.

The SAM and CAM cell bodies are intermingled in the cortex and their primary neurites run together in the same bundles, also previously referred to as the *AM tract* (Gray and Young 1964; Gray 1970; Young 1979). These bundles form a *quasi-columnar* organization toward the neuropil (**Figure 6A, 6B and Figure 7A**). The SAMs and CAMs showed quite differed connectivity, morphology, and ultrastructure (**Figure 6A and 6B**). The SAMs are monopolar interneurons with a single non-bifurcating neurite. Their cell bodies are small (mean ± std; 207.7 ± 26.1 μm^3^; n=69) and contain almost no cytoplasm. In contrast, the CAMs have a complex bifurcating dendritic tree, a larger cell body (mean ± std; 337.2 ± 41.6 μm^3^; n=16) and cytoplasmic volume (**Figure 4**). Unexpectedly, we found that each SAM receives input from only one SFL axon (See SAM section for details). In contrast, each CAM integrates synaptic inputs from a dozen to hundreds of SFL axons and SAMs (**Figure 6B**). Both the CAMs and SAMs consistently converged onto the LN processes (**Figure 7**). Overall, the transmission from the main afferents (SFLs) is entirely mediated via the SAMs and CAMs into their shared target neurons - the dendritic processes of the LNs.

**Figure 4.**
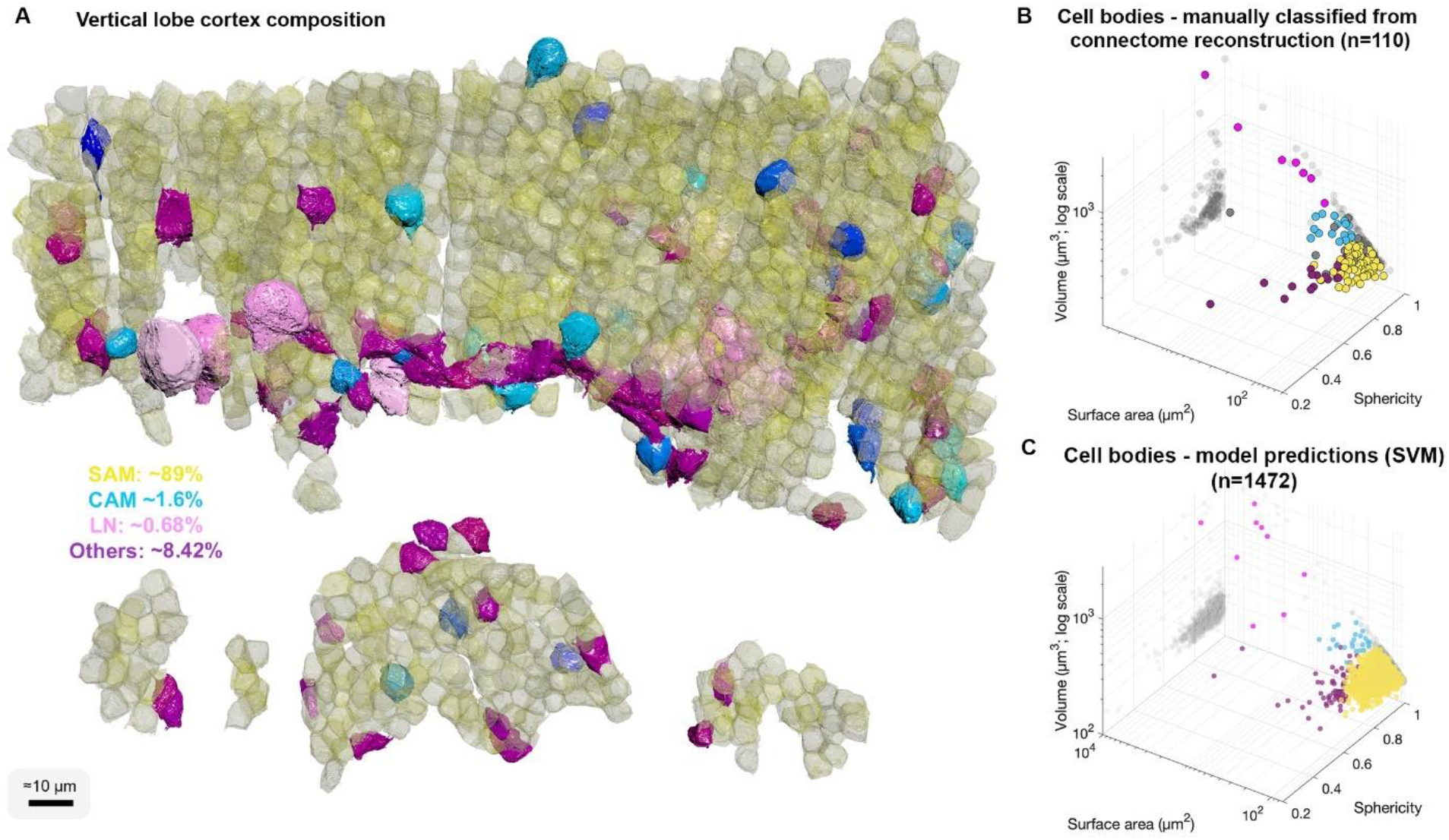
Cell body composition of VL cortex. A. A subset of 1472 densely packed cell bodies were reconstructed and automatically classified based on their morphology; SAM, yellow; CAM, blue hues; LN, pink; Others, purple. B-C. The classifier was designed to capture an observed correlation between cell type and cell body geometry roundish, relatively smaller cell bodies were associated with SAMs (vol 207.7 ± 26μm3, mean ± STD), often larger and slightly more elongated cell bodies with CAMs (vol 337.1671 ± 41.6 μm3), considerably larger cell bodies with highly variable volumes with LNs (volume range 464 – 2850 μm3), and elongated cell bodies with other cell types including glia and uncharacterized cells. A Support Vector Machine (SVM) was trained to separate 4 cell categories SAM, CAM, LN, and Others on a set of 110 manually classified cells based on 3 features: volume, sphericity, and surface area, and was validated on an independently prepared validation set with 47 cells (94.55% training accuracy with ~4.01% error on validation set all associated with SAMs).

A second group of afferent axons innervated exclusively the CAMs. We termed these ‘CAM-input-neurons’ (CIN; **Figure 8**). Even though CIN function has yet to be established, their ultrastructure is reminiscent of the nerve endings which Gray and Young speculated may carry “pain” signals to the VL. They described them as having neurofilaments ending shortly before the synaptic boutons which were filled with small vesicles, a structure we also identified in our data. We also identified that the likely fast transmission circuit (**Figure 2** panels framed with solid line) is overlaid by a diffuse mesh of two types of likely neuromodulatory processes (NM and AF), each with a typical morphology, ultrastructure, and cell-specific connections (**Figure 2**; panels framed with broken line). First, the ‘neuromodulatory fibers’ (NMs) manifest a large variety of processes with large varicosities filled with large vesicles (60-200nm) that often do not show the typical structure of a synaptic connection (detailed below). Second, the ‘ascending fibers’ (AFs) climb from the neuropil up to the cortex through the *AM tracts* (i.e., a dense bundle of primarily SAM and CAM neurites; **Figure 9**). A stereotyped feature of all AFs was their “menorah”-shaped bifurcating pattern, close association with SAM neurites and their intense innervation by the SAM. The ultrastructure and connectivity of these cells revealed two distinct subcategories, one likely neuromodulatory (AF-Nm, **Figure 9C**) and the other yet undefined (AF-Mix, **Figure 9D**). Both the AF-Nm and the NM boutons were filled with large vesicles and usually lacked synaptic outputs, suggesting that these are release sites of neuromodulators or neuropeptides that are involved in extrasynaptic volume transmission (Bentley et al., 2016). Since none of the AF and NM neurites were associated with a reconstructed cell body, we hypothesize that AFs and NMs are external to the VL and may belong to the widespread ascending fibers entering the VL from below that have been previously described (Young, 1971).

Overall, we show that the 7 cell types form a feedforward neural network in which only the AFs display more than 10 reciprocal connections within a cell type (AF-to-AF, **Figure 2**). Analysis at the single neurite level showed that the number of recurrent connections (cycles of connectivity) in the ROI was significantly smaller than that in wiring diagrams built from the same set of neurites and synapses, but with otherwise random directionality determining the pre- and postsynaptic neurons in each connection (p<0.05) (see Methods). Moreover, the connections involved in feedback connectivity *(i.e.*, a cycle of connectivity) were all associated with reciprocal connections which were rare.

### Each SFL axon innervates the SAMs and CAMs via two specialized bouton types

A set of 35 seed SFL afferents was used for a detailed description of the connectivity pattern of the SFL and its postsynaptic targets (**Figure 3A**). Each SFL axon was found to make two distinct types of synaptic contacts depending on the target interneuron. First, an average of 28 ± 12 % (mean ± std) of the synapses of each SFL axon (n=45/161, N=35 SFL) were made on SAMs (**Figure 3B** and **Figure 3–figure supplement 1**). This synaptic connection is unique as each SAM received only a single SFL input. The characteristic synaptic structure consisted of an especially large SFL bouton (LB, 6.41 ± 2.1 μm^3^) innervating *en passant* a specialized vesicle-filled postsynaptic compartment protruding from the SAM primary neurite that partially wrapped the SFL LB (**Figure 3A, Figure 3–figure supplement 1** and **Figure 4**). We termed these postsynaptic structures ‘palms’ due to their somewhat flat and concave morphology. Second, an average of 66± 11% of the synapses (109/161 synapses; N= 35 SFL) of each SFL axon were made on CAM dendritic endings. These CAM endings were a variety of twig-like structures formed by variably long (approximately 1 – 30 μm) thin protrusions emerging from thicker CAM dendritic branches (**Figure 3A**). These postsynaptic profiles were electron-lucent and filled with polyribosomes (**Figure 3B**).

Unlike the single SFL-to-SAM connection, one SFL axon could make several synapses on the same CAM. These SFL-to-CAM synapses showed two different arrangements. Either the synapses involved a large presynaptic bouton (LB) of the SFL manifesting a multisynaptic glomerulus-like structure (**Figure 3A and Figure 5**) or the synapses involved only a single CAM output with a significantly smaller presynaptic SFL bouton (SB, 0.95 ± 0.88 μm^3^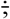 **Figure 3A**). The volume of all boutons (N=295) of all reconstructed SFL axons (N=84) showed a bimodal distribution corresponding to these two different bouton types (**Figure 3C**).

**Figure 5.**
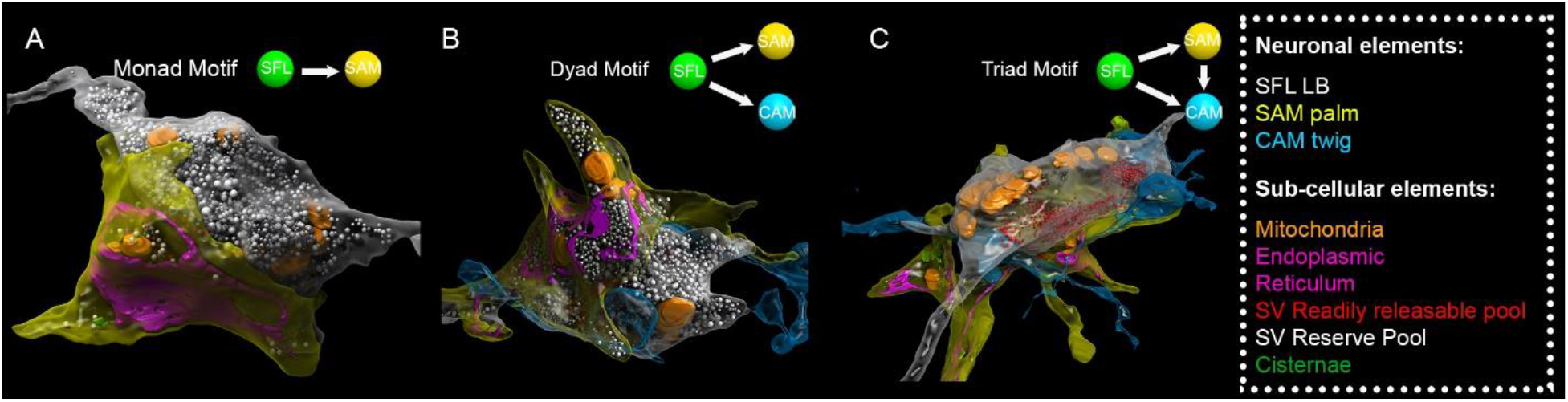
SFL large boutons (LBs) are found in multisynaptic motifs, forming a synaptic glomerulus involving serial connectivity with both SAMs and CAMs. Subcellular 3D reconstruction illustrating the main motifs involving the large varicosity of the SFL axon, which contains glutamate and makes *en passant* synapses. (A) monad (B) dyad and (C) triad. Note that in (C) only the readily releasable pool of synaptic vesicles is shown for clarity as otherwise the large “reserve” pool in the SFL LB (>5000 synaptic vesicles) would obscure other components of the image. The SFL LB were always partly wrapped by at least one SAM palm (yellow) and they often additionally synapsed onto a postsynaptic CAM twig-like ending (blue).

**Figure 6.**
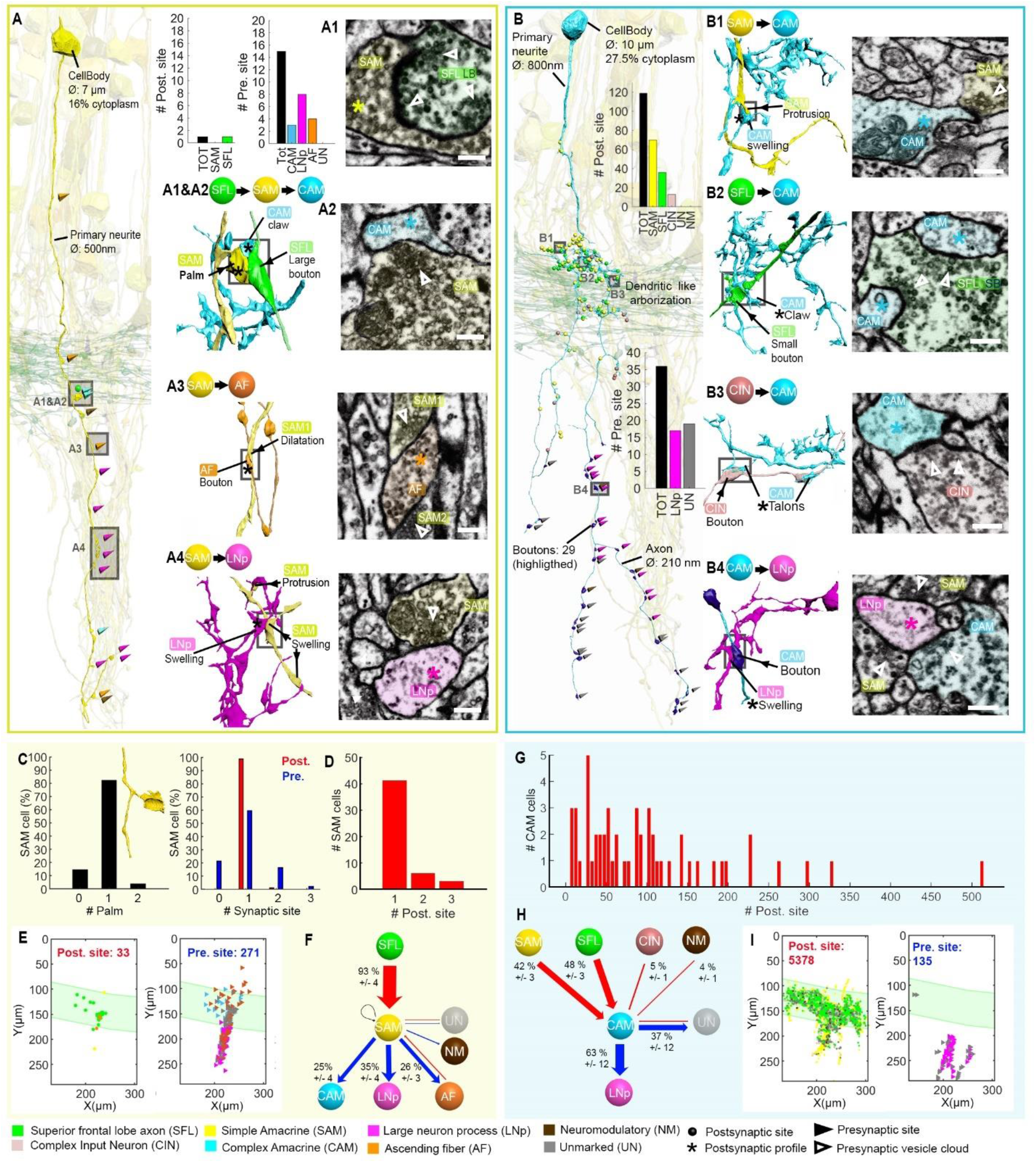
Stereotypic SFL (‘mononeural’)-input to the SAMs and multiple (‘polyneuronal’) inputs to the CAMs. (A,B) 2D projection of a reconstructed SAM (A) and CAM (B), superimposed on a reconstruction of the SFL tract (green). Triangles and puncta represent presynaptic output and postsynaptic input sites, respectively, color coded according to pre- or postsynaptic partner. The insets show the corresponding EM cross-section of each type of synapse. The morphological arrangement of the corresponding synaptic connection is depicted in the colored connectivity scheme next to each image and morphological reconstructions in the corresponding cells’ color code (scale bar=500nm). (C) Analysis of the number of inputs onto the SAM ‘palm’ and its pre- and postsynaptic morphological specialization. 82% of SAMs had only one palm, each receiving just one input (red bar). Each SAM innervated up to 3 targets (blue bars). Only SAMs with their primary neurite fully contained in the SFL tract area were considered (N=56). (D) The majority of the SAMs had only one input located at the palm. (E) Spatial localization of the input/output of the 28 SAM seed cells within the ROI. Note the restriction of SFL-to-SAM inputs to the area of the SFL tract. (F) The connectivity diagram of the 28 SAM seed cells shows the fraction of the inputs (red) and outputs (blue) by neuron type. (G) Histogram showing the wide distribution of the number of synaptic inputs to the CAMs (dozens to hundreds). (H) The connectivity diagram of 53 CAMs shows the percentage of total inputs to and outputs from the corresponding cell type (color coded. See B1,2,3,4 for details). Most of the CAMs’ presynaptic partners were not reconstructed but identified based on characteristic ultrastructure profiles. Profiles which could not be unambiguously identified were classified as unidentified (Un). (I) Spatial distribution of the CAM inputs and outputs within the ROI. See also **Figure 6–figure supplement 1, Figure 6–video supplement 2**, **Figure 6–figure supplement 3** and **Figure 6–figure supplement 4**.

### The number of CAMs is extremely low relative to that of SAMs

We were next interested in the relative abundance the SAMs and CAMs. Since we found significant differences in volume and shape of the various cell types, we reconstructed 1482 cell bodies without their projections and used volume and shape descriptors to build a Support Vector Machine classifier to estimate the number of different types of neurons in the VL (**Figure 4**).

We estimated that 89.3% of the cell bodies in the sampled volume were SAM cell bodies and 1.6% were CAMs (**Figure 4A**) with error rate on validation set smaller than 4.1% (see Methods). Assuming that the VL is roughly composed of 25 million of cells bodies (Young 1963) and a similar proportion over the VL gives an estimate of ~22.3 ± 1.2 million SAMs and an estimate that the cortex contains fewer than 1.43M CAMs. As we did not observe any false negative or false positive associated with CAMs, the total number is likely close to the classifier’s best estimate of 400,000 CAMs which can be further analyzed once this sparse set of cells is sampled on larger EM volumes. Assuming the SFL axons do not bifurcate in the MSF tract, then due to the one-to-one SFL-to-SAM innervation pattern the 1.8 million SFL axons fan-out onto the SAMs at an expansion ratio of about 1-to-12. Since each SAM is activated by a single SFL axon, then information from one SFL axon is represented at the SAM input layer in a non-overlapping manner in only about 12 ± 3.46 SAMs (assuming a Poisson distribution with an average of 12 LB). To estimate the distribution of LBs along the SFL axons we calculated the number of LBs per 100 μm in the 84 reconstructed SFL axonal segments. The results indicate an average and SD of 1.03 ± 0.73 LBs per 100 μm long segments suggesting highly sparse and variable (CV=0.77) distribution along the tract. If this distribution in maintained along large portions of the axon, then the LBs stretch over an average length of 1270 μm which grossly fits the length the of the SFL tract in our preparation (**Figure 1A**). This, in turn, suggests that the LB outputs into the 12 ± 3.46 SAMs is likely maintained along the entire length of the SFL axon rather than being confined in specific area.

### The SAMs, CAMs and SFL large boutons are interconnected within synaptic glomeruli

The SAMs-to-CAMs connection forms the interconnection between the two parallel feedforward networks (**Figure 2**). These connections were found locally at a single large synaptic glomerulus encapsulating the SFL LB, the SAM palm and the CAM twig endings (**Figure 5**). Within this glomerulus these 3 neurons were repeatedly connected with multisynaptic motifs: monad, SFL-to-SAM (**Figure 5A**); dyad, SFL-to-CAM and SFL-to-SAM (**Figure 5B**), serial synapses, SFL-to-SAM-to-CAM, and triad where the same SFL presynaptic terminals innervate an adjacent SAM palm and a CAM twig ending (**Figure 5C**), as well as higher order polyads.

This arrangement fits the classical organization of synaptic glomeruli where a number of synapses are congregated in spatially compact structures, so that they may influence each other’s functioning (Morgan and Lichtman 2020). Among these arrangements, the SAM palm, packed with vesicles, appears to be a unique bifunctional pre- and postsynaptic compartment, as most of the SAM palms formed serial synaptic connections (82%), receiving a synapse from a single SFL bouton and providing a synapse to 1 - 3 postsynaptic compartments, often twig endings of distinct CAMs. Analyzing a large population of ‘long’ SAM neurites (SAMs whose primary neurite was fully contained in the SFL tract area, N=56) revealed that each SAM exhibited only a single palm along its inwardly directed primary neurite (N=46/56), rarely two palms (N=2/56) (**Figure 6C**) and each palm received input from only one SFL large presynaptic bouton (50/50 palms received a single SFL input).

To better describe the structural properties of this unique palm structure we prepared a smaller dataset of 16×16×3.5 μm imaged at 1nm/px (high magnification dataset, see Methods). This revealed the complex presynaptic structure of the SAM palm which could contain up to 1500 synaptic vesicles (36.8 ± 7.7 nm) often tightly packed and resembling the “dense-walled” vesicles described by Gray and Young (1964). The structure included numerous elongated mitochondria and a large network of electron-lucent tubular organelles (**Figure 5**). These organelles could be small, disconnected units (cisternae, Cis) or large and continuous, likely part of the endoplasmic reticulum (ER). The main body of the ER usually faced the pre- and postsynaptic sites, possibly playing a role in vesicle recycling (Ramírez and Couve 2011; Weigel et al. 2021). We also occasionally observed what appeared to be exocytosis at the ER membrane, suggesting the possibility of a Golgi-like extended role in the synaptic vesicle machinery. It is still not clear whether the Cis (up to 15 observed in one palm) were part of the larger ER network or were isolated entities that could, for instance, play a role in keeping vesicles close to the active zone like synaptic ribbons (Dowling et al. 1966; Parsons and Sterling 2003) or the T-bar in Drosophila neurons (Scheffer et al. 2020).

### The SAMs distribute their single SFL input onto the LNs, CAMs and AF

We next focused on a detailed reconstruction and proofreading of 28 SAM seed cells. As mentioned above, each seed SAM neuritic trunk received only one *en passant* SFL LB input (**Figure 6A**). The LB input was always located at the SAM palm (**Figure 5 and Figure 6A1, C**). Analyzing the neuritic trunks of SAM seed cells showed that 93% of the seed SAMs appeared to have no other inputs beside the SFL LB.

The SAM outputs were spread evenly along the length of the neuritic trunk. These synaptic outputs targeted 3 cell types (**Figure 6F**). As described by Young, (1971) and confirmed here, many SAMs converged onto the dendrite-like processes of the LN (LNp), mostly within the central neuropil but occasionally at the level of the SFL tract (**Figure 6A4 and Figure 6E**).

However, the SAM inputs to the LNp represented only 35% of the SAMs’ presynaptic sites (N=109/271). The remaining connections are described here for the first time. 25% of the SAM outputs targeted the CAMs, while 26% innervated the processes of the climbing AFs, suggesting the SAMs directly innervate neuromodulatory processes. The SAM-to-CAM synapses were formed by a variety of dilatations of the SAM neuritic trunk containing clear, small vesicle-filled profiles synapsing onto a clear postsynaptic profile of the CAM twig (**Figure 6A2**). This was mainly observed within the SFL tract and in the upper neuropil. In contrast, the SAM-to-AF synapses (**Figure 6A3**) were spread throughout the dataset, including the cortex, and the postsynaptic element (AF) was a bouton often filled with dense-core vesicles (**Figure 9**). In the neuropil and the SFL tract these synapses were arranged in a rosette-like structure in which 2 to 5 adjacent SAMs synapsed locally onto the same AF bouton (**Figure 9**). Strikingly, within the cortex, the SAM soma occasionally synapsed onto AF varicosities, forming a heterodox somato-neuromodulatory synapse (**Figure 9A**). We also found rare cases of other synaptic outputs to NM (N=2/271), unidentified processes (N=19/271) and only one case of recurrent connection of a SAM onto itself (N=1/271) (**Figure 6F**). Reciprocal synaptic contact between SFL and SAM as well as AF and SAM were occasionally observed.

### Morphology and connectivity of the CAMs

The CAMs were probably not previously identified due to their small number relative to the huge number of SAMs. Gray (1970) and Young (1971) may therefore have confused their dendritic arborizations and electron-lucent ultrastructure within the SFL tract (**Figure 6A2,B1,B2 and B3**) with that of the LN. To understand the potential functionality of these newly discovered interneurons within the VL circuitry we annotated all their pre- and postsynaptic sites. Because the SFL, CINs, SAMs AFs and NMs all have distinctive ultrastructure “fingerprints” *(e.g.*, cytoplasm staining, vesicle size, vesicle type, mitochondria shape; see Methods), we were able to define their presynaptic partners without reconstruction.

The CAM neurites have 3 distinct regions: the *primary neurite* ramifies into strictly postsynaptic *dendritic branches* in the SFL tract. The dendritic branches turn into neuritic *axons* in the deeper neuropil (**Figure 6B and Figure 6-figure supplement 2**). Synapses were located preferentially on numerous small twigs which branched off the thicker dendritic branches. We used ‘twig’ to define CAM second-order dendritic branches because their base lies on a microtubule-containing main thick branch(Schneider-Mizell et al. 2016) but, unlike spines that have a maximal length of less than 3 μm (Leiss et al. 2009), the twigs can be much longer and show a complicated branching pattern. The CAMs showed variable dendritic branch patterns and orientation, possibly suggesting the existence of CAM subtypes (**Figure 6-figure supplement 2 and 3**). Additionally, the morphology of the CAM twig endings was highly variable with at least 5 categories all somewhat resembling dendritic spine enlargements (**Figure 6B,C**): flatfoot, long neck mushroom head, talon, palm-like and a claw-like endings that strikingly resembled the claw-like structure wrapping the projection neurons in the insect MB (Leiss et al., 2009; Schürmann, 2016; Yasuyama et al., 2002).

In contrast to the SAMs (**Figure 6–figure supplement 1**), each CAM interneuron received dozens to hundreds of synaptic inputs (**Figure 6G and Figure 6–figure supplement 2 and 3**); the most extreme case displayed 514 postsynaptic sites over a neurite length of 1.4μm. The CAMs integrate a combination of inputs (**Figure 6H**), mostly from the SFL (48%, **Figure 6B2 and Figure 6–figure supplement 2G and 3E**) and the SAMs (42%, **Figure 6B1 and Figure 6–figure supplement 2F and 3F**). 5% of the inputs to the CAMs originated from CINs (**Figure 6B3 and Figure 6H**). The CINs appeared to exclusively innervate the CAMs with their inputs sparsely distributed over the CAM twig endings (**Figure 8**). Despite the CAMs showing high diversity in primary neurite projection and dendritic arborizations, all received inputs and innervated the same cell types and at similar locations within the VL (**Figure 6B and Figure 6–figure supplement 3 and 4**). The CAMs’ distal axonal processes innervated the LNps with many *en passant* boutons (**Figure 6B4**).

### Different convergence patterns of SAMs and CAMs into the LNs

Both the SAMs and CAMs converged onto the LNs, the output cells layer of the VL (**Figure 2, Figure 6 and Figure 7**). To investigate the pattern of LN innervation we first annotated all the synapses of 14 LNs with reconstructed cell bodies and those on putative LN processes (see below, N=24). The LN cell bodies (dia. > 15 μm) and their radially directed large primary neurite were poorly contained within our analyzed volume. However, some of them (N=3/14) gave off dendritic collaterals within the SFL tract area where they were solely postsynaptic to SAMs (**Figure 2**) and to other unidentified processes (with a total of 115 individual synaptic inputs on the LN processes in the SFL tract).

**Figure 7.**
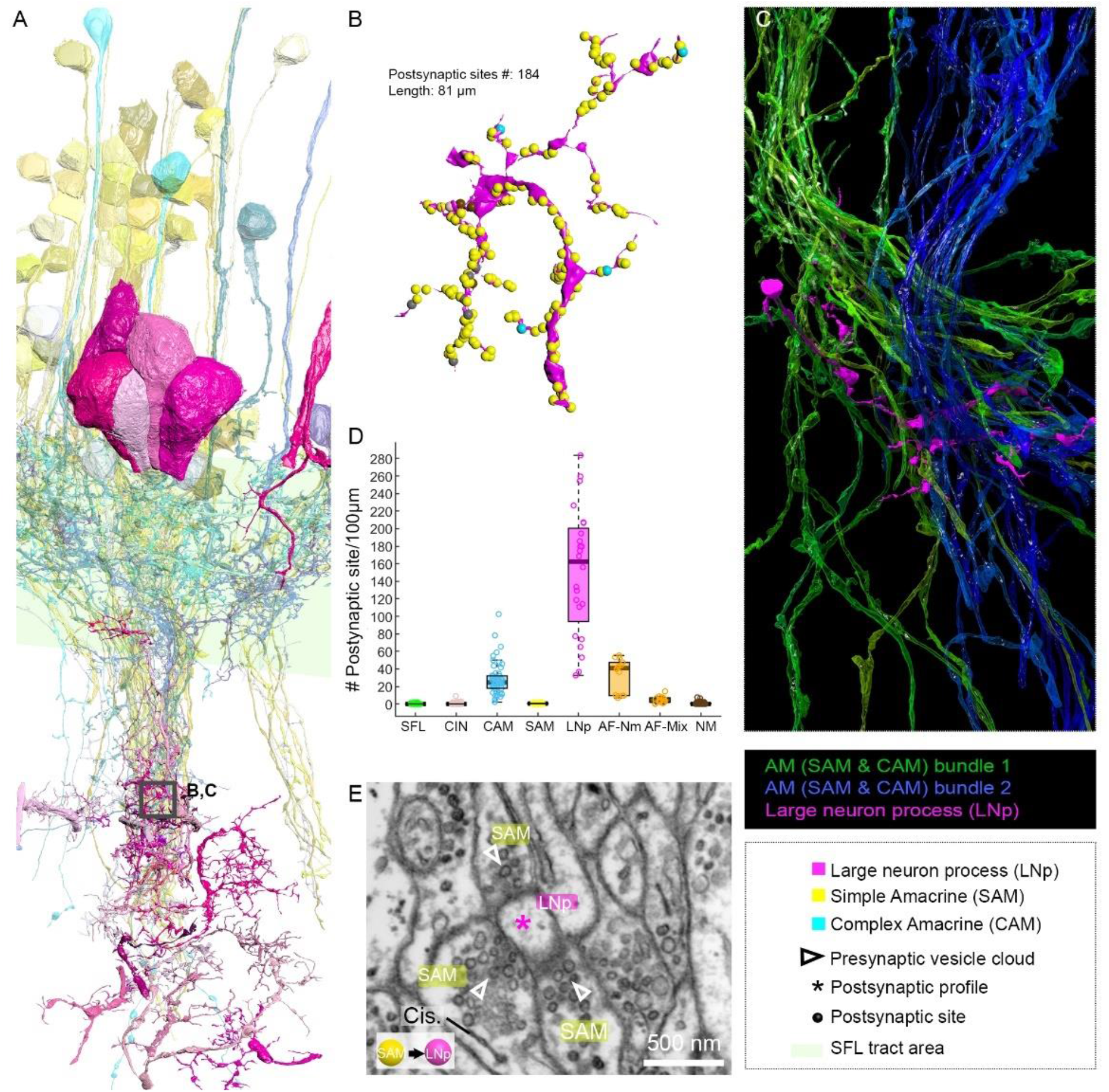
Patterns of synaptic convergence of SAMs and CAMs onto LNs. (A) 3D reconstruction of the LN cell bodies, and processes (LNp) (magenta hues) superimposed on SAMs (yellow) and CAMs (cyan). (B) Distributions of the 184 postsynaptic sites on one LN dendrite colored according to the legend. Note the very few CAM inputs relative to those of the SAMs. (C) Convergence of two bundles of intermingled SAM and CAM neurites onto an LNp shown in B. (D) The LNps displayed the greatest postsynaptic density among all dendritic elements. (E) High resolution EM illustrating a rosette-like structure formed by the convergence of 3 SAMs into one LNp. Empty triangle and asterisk are pre- and postsynaptic profiles, respectively. Resolution: 1nm/px. Dwell time: 200 μs.

**Figure 8.**
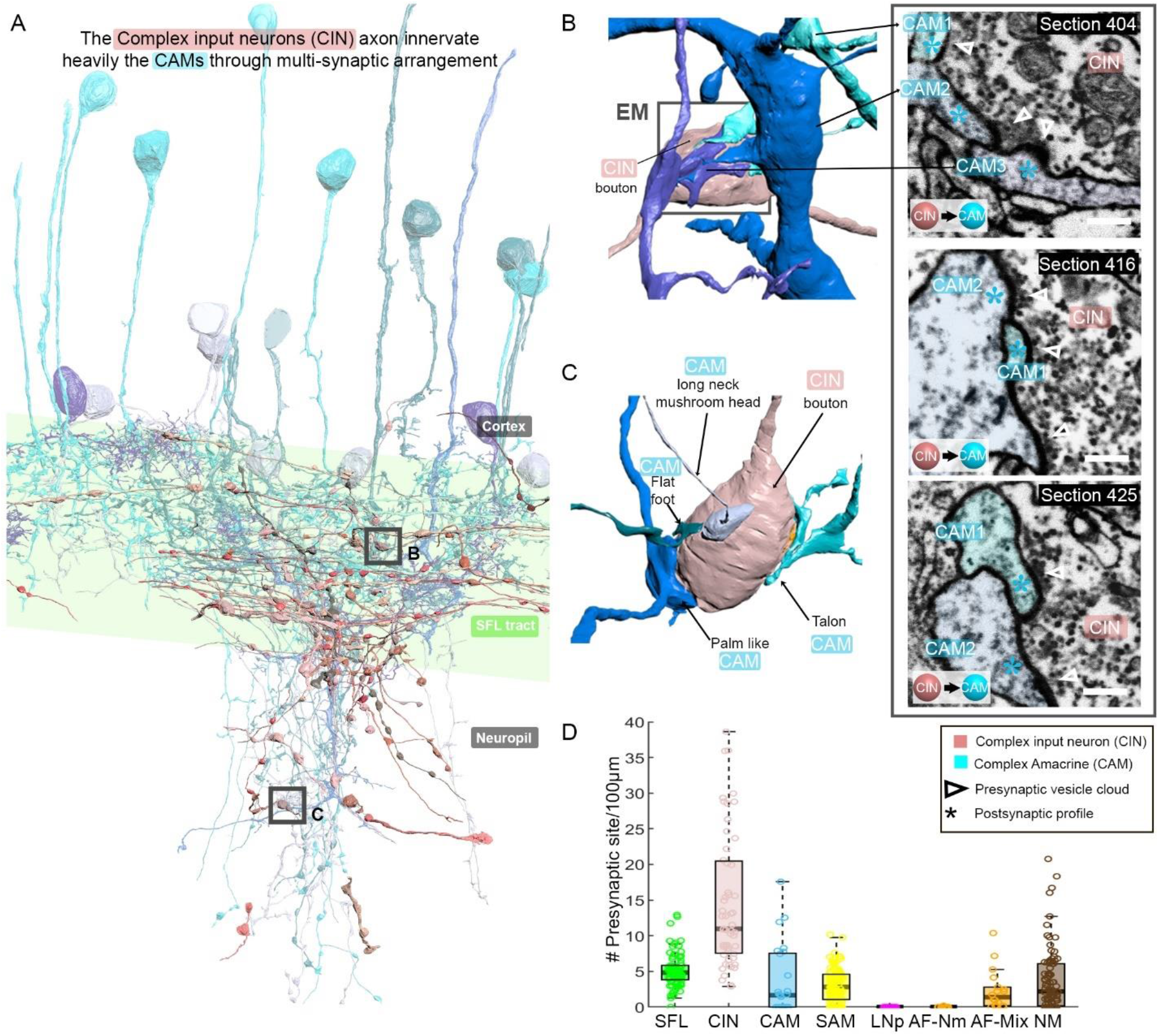
CAM-specific afferents (CINs) innervate exclusively CAMs via large multisynaptic boutons. (A) 3D reconstruction of a CIN (pink) superimposed on the CAMs, their main postsynaptic partners (blue hues). (B, C) 3D arrangement of a large CIN presynaptic bouton innervating multiple CAM dendrites (3 in B; 4 in C). CAM twig endings varied in shape resembling flattened dendritic spine enlargements. As indicated in C their structure can be divided into at least 5 classes. Right inset, 3 consecutives near cross-sections through a CIN bouton and its 3 postsynaptic targets. The CIN profile contained the smallest vesicles, about 30nm dia., of the entire dataset. (D) CIN had the highest presynaptic site density of the dataset. Error bar=SEM.

**Figure 9.**
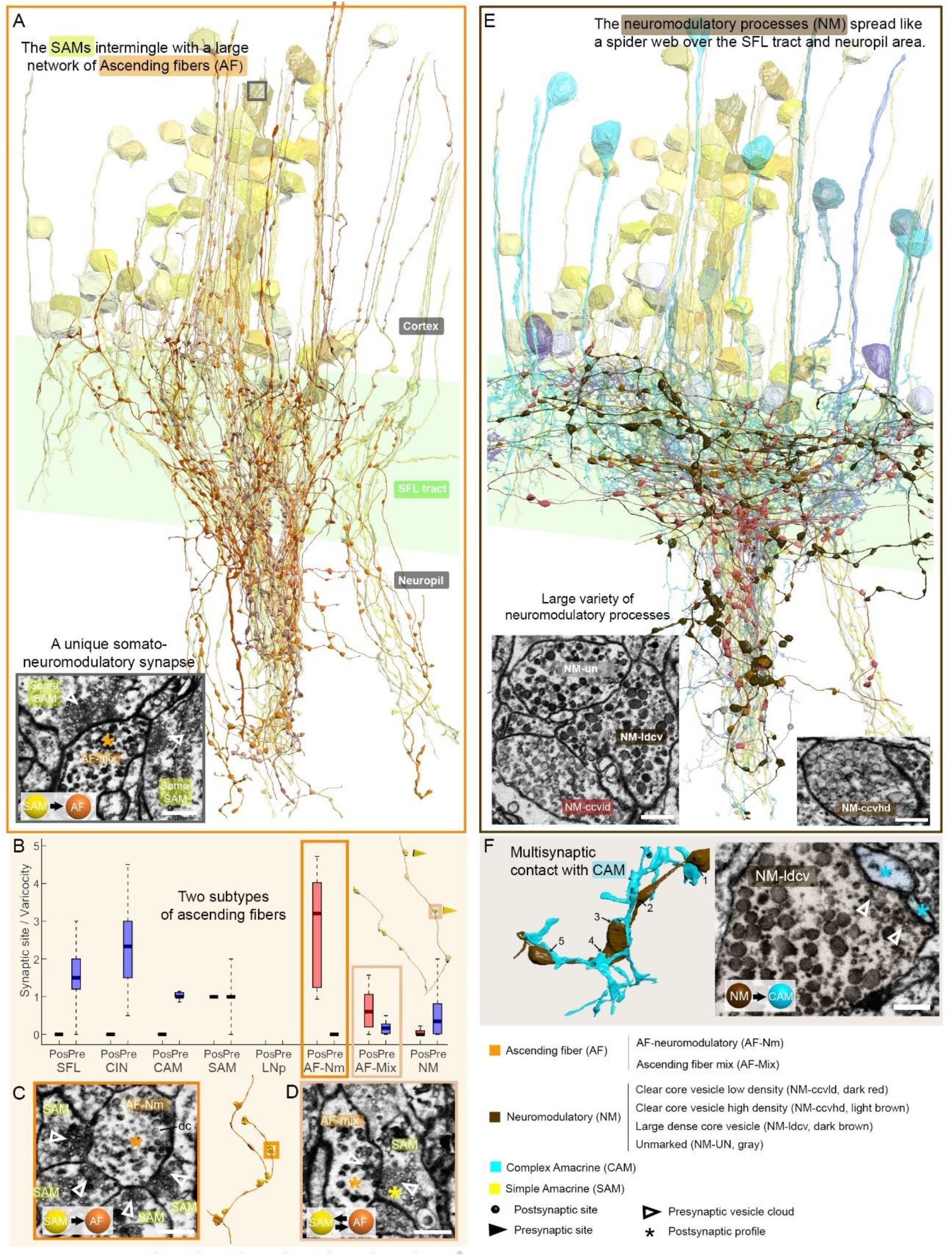
Overlay of two modulatory systems, the ascending fibers (AFs) and widespread neuromodulatory fibers (NMs). (A) 3D reconstruction of a large network of AFs (orange) intermingling with a SAM neuritic bundle (yellow). The AFs showed varicosities along their length filled with large vesicles and were mainly postsynaptic to the SAMs (C,D) A somatic synaptic input from SAM to AF was occasionally observed (lower left inset). (B) The distribution of the number of pre- and postsynaptic sites per varicosity (blue and red, respectively) revealed two types of AF processes shown in C and D. (C) The AF-Nm type (orange) were exclusively unidirectional synapses forming a rosette-like structure where 2-5 SAMs made adjacent synapses onto one AF-Nm postsynaptic varicosity (asterisk). (D) The AF-mix (light orange) were more diverse. Their boutons were filled with vesicles of various shapes and sizes, ranging from small dark core vesicles to flat clear core vesicles. Unlike the AF-Nm, they could be bidirectional, both pre- and postsynaptic (triangles and puncta, respectively) and occasionally displayed reciprocal contact with a SAM as shown in the EM. (E) 3D reconstruction of the widespread NM fibers color coded by subtype and superimposed on populations of SAMs (yellow) and CAMs (blue). At least 3 distinct subtypes were distinguishable based on vesicle sizes (EM insets: clear core low density, clear core high density, dense core). (F) 3D reconstruction of NM-to-CAM synapses. The NM made multisynaptic contacts. The right inset shows a representative EM of a neuromodulatory synapse. Scale bar EM sections, 500 nm. See **Figure 9–figure supplement 1**

As suggested by the physiological experiments (Shomrat et al. 2011), we did not find any direct connections between these LN dendritic collaterals and SFL axons. Among the 7 cell types thoroughly studied, we consistently observed that the SFL, SAMs, CAMs, CINs, AFs and NMs processes all displayed an EM profile filled with vesicles within the neuropil. Therefore, by elimination, we considered all electron-lucent postsynaptic profiles within the neuropil as putative LN dendritic processes (LNp). The 24 LNps that we examined branched profusely within the neuropil (**Figure 7A**) and were heavily innervated. The histogram in **Figure 7D** shows that the LNps displayed the largest density of incoming synapses in the dataset with a median of 162 postsynaptic sites/100 μm, far larger than the densely innervated AF-Nms (41 postsynaptic sites/100 μm) and the CAMs (25 postsynaptic sites/100 μm that are restricted to the SFL tract; **Figure 7D**). The SAM and CAM innervation of the LNps often formed rosette-like structures in which 2 - 5 or more AMs made adjacent synapses onto an LN dendritic process (**Figure 7E**).

We next estimated the degree of convergence of the SAMs and CAMs onto the LNs. For this we reconstructed and annotated all the neurites synapsing onto an 81 μm length of LNp dendrite (**Figure 7 B, C**). As shown in **Figure 7C** all the CAM and SAM inputs originated from two distinct fasciculate neuritic bundles (blue and green) that included 160 SAM neurites and only six CAMs. Approximately 90% of the inputs on LNps were from SAMs (N=160/184 postsynaptic sites), only 3% were from CAMs (n=6/184 postsynaptic sites). **Figure 6B** shows that the CAM and SAM inputs were distributed in a “salt and pepper” manner with the CAM inputs distributed very sparsely and evenly over the LN dendritic process. These data indicate that the VL output neurons integrate massive information from the SAMs and only sparse input from the CAMs. The spatial distribution of the CAM inputs to the LNp suggests a role of the CAM output in adjusting the membrane potential of a large LN dendrite area. Such spatial distribution would be most effective if the CAMs are inhibitory interneurons as we suggest below (see Discussion).

### The CINs afferents innervate exclusively CAMs

CIN processes spread throughout the neuropil, but many ran parallel to the SFL axons (**Figure 8A**). Their boutons were filled with many small, rounded mitochondria and the smallest vesicles observed in the VL (≈30nm). Their presynaptic boutons displayed the greatest variability in size with volumes ranging from 0.09 to 29.50 μm^3^ (median ± iqr; 1.65 ± 2.23).

Fifty-five of these fibers were reconstructed and their synaptic sites annotated. The CINs were almost exclusively presynaptic (534 sites) with only very few postsynaptic sites (17 sites). All these presynaptic sites were located on large boutons (N=239) that usually made polyadic synapses with several CAMs’ postsynaptic structures *(e.g.*, 3 in **Figure 8B** and 4 in **Figure 8C**) with a median of 2.3 ± 1.5 (median ± iqr) synaptic sites per bouton (**Figure 8D**), the most extreme case being 8 presynaptic sites for a single bouton. This synaptic structure may allow a single CIN terminal to influence several CAMs simultaneously.

The CIN innervate *en passant* mainly the CAMs (225 synaptic connections) and only occasionally the AFs (**Figure 2 and Figure 2–figure supplement 1**). Note that the CAMs projected long thin side branches with a morphological specialization at the tip *(i.e.*, long neck with mushroom head, **Figure 8C**) where it received a synaptic input from the CIN bouton. The CINs displayed the highest presynaptic site density found within the VL with a median of 11 presynaptic sites/100μm, twice as high as presynaptic density on SFL axons (**Figure 8D**).

### The neuromodulatory ascending fibers (AF) receive massive synaptic inputs from the SAMs

The AF were highly recognizable in our dataset with their menorah shape, climbing through the neuropil into the cortex while intermingling with SAM and CAM neurites (**Figure 9A**). Their processes displayed large boutons along their length (9.4 boutons/100μm). The reconstruction of the synaptic partners of the SAM seed cells revealed that the AFs receive massive input from the SAMs (**Figure 6F**).

Mapping the AF synaptic sites revealed two clear subcategories of AF based on their connectivity (**Figure 9B**) and EM profile (**Figure 9C,D**). We termed one class AF-neuromodulatory (AF-Nm, n=14). In this class, on the 468 boutons reconstructed we annotated 1460 postsynaptic sites and only 3 presynaptic sites, with a median of 3.2 postsynaptic site per bouton. Each bouton formed a rosette structure of axo-axonal synapses where several SAMs made adjacent synapses onto one AF bouton (**Figure 9C**). The postsynaptic sites were filled mostly with large dense core vesicles (60-80 nm), suggesting a neuromodulatory nature. The synaptic inputs of the SAMs into the AF-Nm boutons may suggest SAMs-induced local extrasynaptic release of neuromodulator (see Discussion). The AFs seem morphologically equivalent to the serotonergic processes ascending into the cell cortex found in immunohistological studies (Shomrat et al. 2010; Shigeno and Ragsdale 2015) and also to tyrosine hydroxylase (catecholaminergic) labeled processes (Stern-Mentch et al., 2022).

In contrast, the second class of AF, the AF-mix (N=18), were more diverse with boutons filled with vesicles of varying shape and size (30-90 nm; N=519), including small dark core vesicles, flat clear core vesicles and larger vesicles. Unlike the AF-Nm, the AF-mix were both presynaptic (median of 0.2 presynaptic sites/bouton) and postsynaptic (median of 0.6 postsynaptic sites/bouton) and occasionally displayed reciprocal contact with a SAM (**Figure 9D**). The diverse nature of the synaptic vesicles in the two subtypes suggests that they convey different neuromodulatory signals and the reciprocal connection with the SAMs suggest their involvement in SAM -induced local release of neuromodulators.

### The ubiquitous NM processes appear to be involved in global neuromodulatory signaling via extrasynaptic transmission

A large variety of processes displayed the largest dense vesicles (60-200nm), which appear similar to vesicles thought to be neuromodulatory in other invertebrates (Messenger 1996; Witvliet et al. 2021). The processes were widespread within the neuropil and the SFL tract and their lack of specific orientation made it impossible to assume their origin (**Figure 9E**). At least 3 distinct profiles were clearly distinguishable by vesicle size and appearance, showing either a low density of clear core vesicles, a high density of clear core vesicles or very large dense core vesicles. The boutons of each process contained only one type of vesicle, which were also present within the fiber. Reconstructions of NM processes (N=88) showed that most of the boutons had neither inputs nor outputs (464/632), suggesting that they were neuromodulatory afferents which exert their effects through extrasynaptic volume transmission. Yet, these cells occasionally synapsed with a variety of postsynaptic partners including CAMs, SAMs AFs, NMs themselves, and LNs (**Figure 2 and Figure 2 - figure supplement 1**). Multisynaptic contacts were occasionally observed, for instance, with CAMs (**Figure 9F**), but the density of synaptic sites per bouton was rather low, 52 presynaptic sites and 250 postsynaptic sites on 632 boutons. Accordingly, NM processes may be involved primarily in extrasynaptic transmission and thus may convey a global modulatory signal transmitted though action potentials propagating along them.

## DISCUSSION

The novel connectivity and anatomical features discovered here profoundly advance our understanding of the functional organization of the learning and memory circuits in the vertical lobe (VL) of *Octopus vulgaris*. The EM volume reconstruction revealed several novel features that, on one hand, show the VL is composed of several more components than previously thought. On the other hand, these novel cell types and their connectivity help us understand the functional organization of the VL as a learning and memory acquisition network.

### The amacrine interneurons in the VL input layer are not a homogeneous group

The first striking finding was that 89% of the ~25 million VL cells (**Figure 4**) send a simple non-bifurcating neurite into the neuropil and were therefore termed simple AMs (SAMs; **Figure 6A**). The SAMs are highly input specific as each received *en passant* input from only a single SFL axon via large presynaptic boutons (LB) (**Figure 3A, Figure 5 and Figure 6A1**). If the pattern of one-to-one innervation is maintained throughout the whole SFL tract and if, as generally believed and as found here, the SFL axons bifurcate very rarely, then every SFL neuron is represented by only around 12 ± 3.46 SAMs (22×10^6^ SAMs / 1.8×10^6^ SFLs). Moreover, the LBs seem to be sparsely distributed along the SFL axon (1.03 ± 0.73 LBs per 100 μm; see Results), suggesting that each SFL neuron is represented in the SAMs in a highly sparse and non-overlapping fashion.

This extremely low and random topographical expansion of SFL inputs to SAMs is unique for input circuitry. In *Drosophila* mushroom bodies, for example, the expansion ratio is multiplied by a factor of about 6 as six projecting neurons converge onto each Kenyon cell (Li et al. 2020) and in locust the expansion ratio is 400 (Jortner et al. 2007). The unique low expansion of information in the VL is achieved by the formation of a huge number of SAMs, 3-4 orders of magnitude more than the number of the homologous Kenyon cells in the insect mushroom bodies. This mode of expansion may be related to the organization of the octopus visual system where each SFL axon carries visual information that has been highly processed by the 120×10^6^ neurons of the two optic lobes and then possibly further classified into lower dimensional visual features in the only 1.8×10^6^ cells in the SFL (Young 1971). This unusual expansion pattern resembles the 50×10^12^ granular cells in the human and mice cerebellum - each receiving only 4-6 inputs from the mossy fibers at the expansion ratio of 1-to-200 (D’Angelo 2013; Kawato et al. 2021). An additional striking similarity is that these unique synaptic connections in the cerebellum undergo activity-dependent presynaptic LTP; (Mapelli and D’Angelo 2007) like the SFL-to-SAM synapse in the octopus VL.

Our second novel discovery is that the AMs included a small group of a new class of AM neuron, only 1.6% of the AMs in the VL cortex (**Figure 4**). These neurons showed a bifurcating dendritic structure and were therefore termed complex AMs (CAMs, **Figure 6B**). The primary neurite segments of the SAMs and CAMs travel together in the AM bundles within the VL cortex, but the non-bifurcating SAM neuritic trunks cross the SFL tract to pass into the inner neuropil, while the CAM neurites bifurcate substantially within the SFL tract (**Figure 2 and Figure 6B**). The CAMs receive inputs *en passant* from multiple SFL axons either directly or indirectly via a serial connection from the SAMs (**Figure 5 and Figure 6B**). Thus, while each SAM represents a single SFL neuron, the CAMs integrate information from dozens to hundreds of SFL neurons together with multiple inputs from nearby SAMs. In addition, as discussed below, the CAMs receive inputs from afferents from a yet unidentified origin, which exclusively innervate CAMs (CIN, **Figure 6B3 and Figure 8**).

### The SFL axons innervate the SAMs and CAMs via different classes of presynaptic boutons

SAMs and CAMs are innervated by SFL axons via small (SFL SB) or extremely large boutons (SFL LB – **Figure 3**). The LBs are always associated with at least one SAM palm and may additionally innervate a CAM twig within a dyadic or triadic arrangement (**Figure 5**). By contrast, SBs are always associated with a single postsynaptic CAM compartment (**Figure 3**). This suggests that the two different types of SFL axon terminal are functionally specialized. Such functional segregation is also seen in the hippocampal mossy fiber connections where small boutons target GABAergic interneurons, whereas large boutons contact CA3 pyramidal cells and hilar mossy cells (Acsády et al. 1998). The latter large synapse shares several synaptic features with the octopus VL synapses (Hochner et al. 2003) – NMDA-independent, presynaptically expressed LTP and an extremely large range of presynaptic short-and long-term synaptic gain modulations (Nicoll and Schmitz 2005; Henze et al. 2000; Hainmueller and Bartos 2020; Restivo et al. 2015). The large synaptic boutons here, like the mossy fiber terminals, may provide an increased dynamical range of synaptic gain control needed for certain forms of plasticity involving presynaptic modulation of transmitter release (Orlando et al. 2021; Hainmueller and Bartos 2020; Restivo et al. 2015).

### The single SFL synaptic input into the SAM may serve as a gain control site of the stimulus-specific input to the VL

Each SAM receives synaptic input from only a single SFL neuron, raising the question of the role of such apparently non-integrating interneurons. Physiological results indicate that the glutamatergic SFL-AM synapse is the site of activity-dependent LTP in the VL (Hochner et al. 2003; Shomrat et al. 2011; Turchetti-Maia et al. 2018). The large volume of the SFL-LB relative to that of the SFL-SB found here (on average 6-fold larger) suggests that the recorded synaptic field potential (fPSP) mainly reflects the synaptic input from the SFL LB to the SAMs. The morphological dimensions of the SAMs revealed here *(e.g.*, a 166μm long 0.5μm diameter neuritic trunk emerging from a 7μm diameter cell body) together with previous electrophysiological measurements (>1GΩ input resistance measured at the cell body), suggest that the inexcitable SAMs (Hochner et al. 2003, 2006)are electrotonically compact (estimated length constant λ > 147 μm) (Hochner and Spira 1987), so that the glutaminergic excitatory postsynaptic potential from the SFL LB input site is likely sufficient to activate voltage-dependent transmitter release down the SAM neurite without needing a propagating action potential. These inexcitable properties, which differ drastically from the excitable properties of the insects Kenyon cells (e.g. Demmer and Kloppenburg 2009), explain nicely the linear input/output relationship of the SAMs (Shomrat et al. 2011). Since the activity-dependent changes at the SFL-SAM synapse involve pre- and postsynaptic mechanisms (see below), we suggest that, rather than being simple relay interneurons, the SAMs provide a synaptic “gain-control relay” of the SFL-LB input into the CAMs and LNs.

### How can specificity and associativity be achieved within these one-to-one SFL-to-SAM connections?

*Specificity* and *associativity* are essential properties for all associative learning networks (Kandel et al., 2012; Malenka 2003). To understand how they are achieved within the unique wiring of the VL it is important to recall that LTP expression in the VL involves presynaptic facilitation of transmitter release from the SFL boutons. This was suggested to be mediated by an activity-dependent persistence elevation of NO concentration in the vicinity of the SFL-SAM connection. NO alone cannot induce LTP. Only the association of strong activation of an SFL-SAM synapse switches ON the NO-dependent NOS activation, leading to long-term LTP *maintenance* (Turchetti-Maia et al. 2018). The good correspondence of the SAMs’ morphological characteristics with AM-specific labeling by NADPH-diaphorase and cChAT (choline acetyl transferase) suggests that NOS activity is strongly expressed along the entire length of the cholinergic SAMs (Stern-Mentch et al. 2022). Accordingly, NO, which can readily diffuse short distances through cell membranes (Agnati et al., 1995; Garthwaite, 2016, 2020; Turchetti-Maia et al., 2017, 2018) and which is generated in the SAMs following LTP induction, diffuses retrogradely from the SAM palm into the presynaptic SFL LB mediating the NO-dependent presynaptic *expression* of LTP. The characteristic morphological arrangement described here with the SAM palm partly wrapping the SFL LB, ensures synaptic specificity as the steep NO concertation gradient will affect only these closely attached presynaptic varicosities (Hall and Garthwaite 2009). The SFL-SAM synapses are organized in a compact synaptic glomerulus that may facilitate the diffusion of NO within a triadic synaptic organization including the CAMs. This suggests that NO diffusing from a single postsynaptic SAM palm also modulates the SFL SB input into the CAMs.

What are the possible mechanisms of LTP *associativity?* The persistence elevation of NO concentration in the VL is maintained, at least in about half of SFL-SAM synapses, through a molecular switch consisting of a positive feedback loop of NO-dependent reactivation of NOS (Turchetti-Maia et al. 2018). This “molecular switch” provides a tentative molecular model for associativity, whereby volume-type transmission of NO (Garthwaite 2016) between adjacent activated SAMs may mutually facilitate associative switching ON of NOS in these simultaneously activated SAMs. Our results highlighted that many SAM neurites converging within the neuropil to form compact bundles of thin intermingling processes travelling together (**Figure 7A**). Such spatially closed arrangements of thin neurites (e.g., **Figure 7A, C**), which express intense NOS activity (see Turchetti-Maia et al., 2018, Stern-Mentch et al 2022), can favor mutual NO effects among adjacent SAM neurites.

### The CAMs as a putative feedforward inhibitory network balancing the parallel SAM feedforward excitatory network

The newly discovered CAMs comprise only 1.6% of the VL cortex. An inhibitory nature of CAMs is supported by GABA labeling of putative CAM dendritic processes at the level of the SFL tract and in varicosities in the neuropil (Stern-Mentch et al. 2022). Also, inhibitory CAMs would explain the IPSPs or EPSP followed with short delay by an IPSP recorded in some of the LNs in response to SFL tract stimulation (**Figure 7 - figure supplement 1**).

Although the CAMs show morphological diversity (**Figure 6-figure supplement 2 and 3**), they share similar connectivity, integrating multiple inputs from the SFLs, SAMs and CAM-specific inputs (CIN) in their bifurcating dendrites (**Figure 6B**). The CAMs innervate the LN dendritic processes only sparsely, unlike the dense LN innervation by the SAMs (**Figure 7B and 7D**). This innervation pattern suggests that the LNs integrate stimulus-specific excitatory inputs from many SAMs, while the CAMs, which integrate ubiquitous and nonselective inputs, are optimally located for inhibitory balancing the overall SAM excitatory inputs. Yet, the LNp are innervated by only a few CAMs, suggesting that the CAMs inhibitory inputs are activated simultaneously over a large LN dendritic area causing widespread postsynaptic inhibition of the LNp.

If the CAMs are indeed inhibitory interneurons, then our results strongly suggest the VL network is organized into two interconnected parallel feedforward networks, in which the SFL axons conveying sensory inputs ‘fan-out’ onto SAMs (in ~1/12 ratio) and ‘fan-in’ (~4/1) onto the CAMs. Both the SAMs and CAMs feedforward onto the LNs in the VL readout layer (schematically shown in **Figure 10A**). This type of organization may sharpen the stimulus specific SAM inputs to the LNs by subtracting the global or background activity monitored by the inhibitory CAMs. This type of organization may highlight a universal inhibitory balancing organization in networks with different functions. A similar organization is found in cortical circuits representing sparse, odor-specific, activity where the sparse activity is generated through summing unbalanced widespread and broadly tuned inhibition with odor-specific excitation (Poo and Isaacson 2009). In the octopus VL, such sharpening mechanism may ensure recognition of learned visual scenarios even under variable light intensities. Theoretical studies have demonstrated the importance of implementing feedforward inhibition parallel to the feedforward excitation to balance the stable propagation of an activity wave (Litvak et al., 2003). Feedforward inhibition also appears important for balancing networks’ input/output relationships to avoid attraction of network activity into either up or down states (Ferrante et al., 2009). Finally, long-term balancing inhibition is believed to be important in ensuring network homeostasis especially in networks involved in learning and memory, like the hippocampus, where activity balances may be perturbated by long-term activity-dependent plasticity (Turrigiano and Nelson 2004).

**Figure 10.**
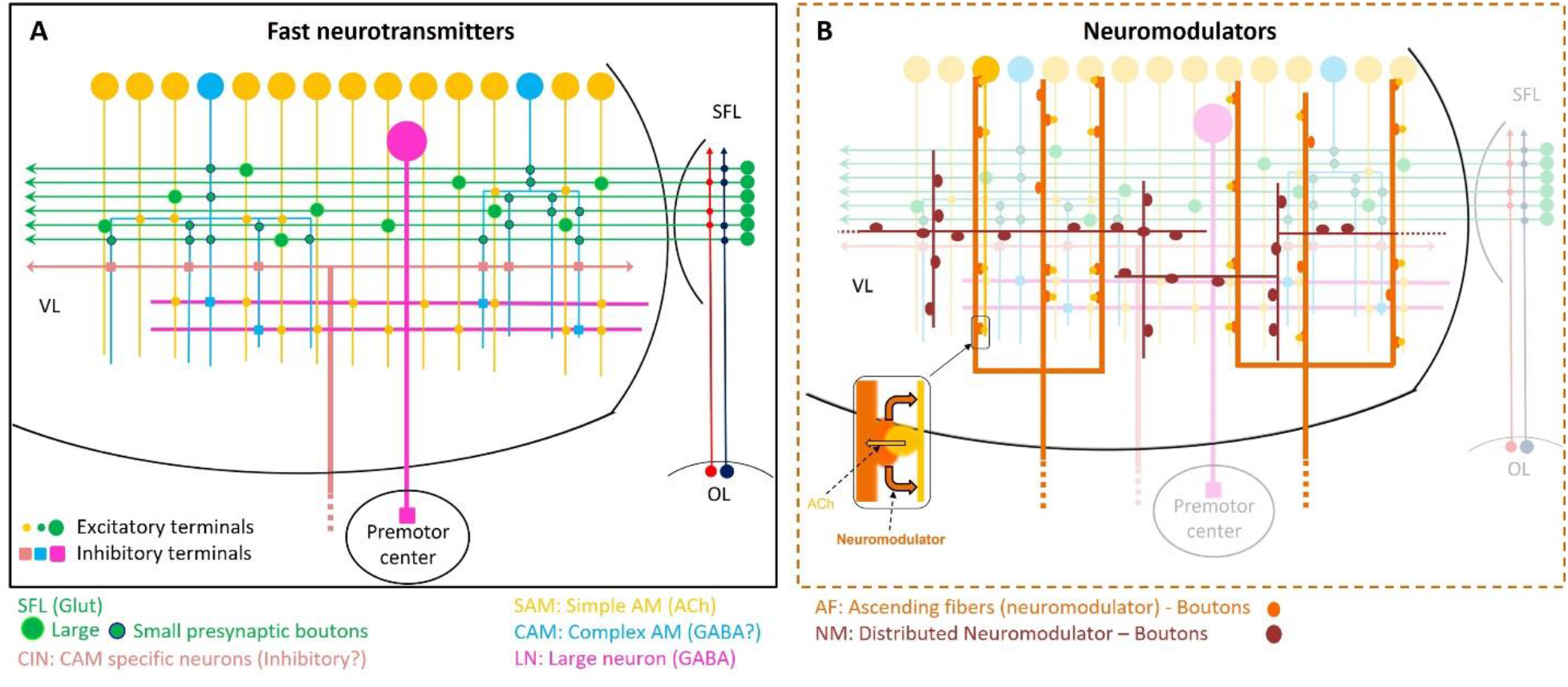
Schematic representation of the currently known circuit architecture of the VL. (A) Fast transmitter connectivity. The parallel SFL axonal tract (green) conveys classified sensory information via *en passant* glutamatergic (glut) synapses to a large group of SAMs (yellow) and a very small group of CAMs (cyan). Each SAM has one palm on its neurite innervated by a single large SFL bouton (SFL LB). This connection is endowed with short- and long-term plasticity. The CAMs integrate ongoing activity through multiple inputs from the SFL, SAMs and CINs (light pink). The SAMs and CAMs converge onto the LN processes (LNp, magenta) forming two parallel and interconnected feedforward networks (shown in **Figure 11**). SAM output to the LNs is excitatory cholinergic (ACh). The CAMs inputs to the LN may be inhibitory GABAergic (see text). (B) The two widespread networks of modulatory inputs into the fast transmission circuit (shown in A). The ascending AF processes (orange) make reciprocal connections with the SAM neurites, possibly inducing local release of neuromodulator (likely serotonin). The NMs (brown) spread throughout the neuropil. Physiological and immunohistochemical studies indicate that AFs and NMs release neuromodulators such as serotonin and octopamine and dopamine that likely play a role in reinforcing/suppressing the LTP at the SFL-to-SAM synapse.

### Three candidates for conveying modulatory signals

Learning and memory networks are usually supervised by neuromodulation. Early anatomical studies by Young and colleagues proposed several candidates, and our study has refined Young’s broad classification of fibers ascending to the VL. We have described 3 distinct classes of putative extrinsic processes: neuromodulatory fibers (NM), ascending fibers (AF) and CAM-specific input fibers (CIN). These could play a role in conveying information on whether the current environment is punitive or rewarding, reinforcing, or suppressing associative learning.

The rich and widely distributed NMs intermingle with the fast transmitter feedforward network (schematized in **Figure 10B**). The fibers form a mesh in the neuropil and may be associated with regulation of overall VL activity. Their exceptionally large boutons, frequently lacking the typical release sites, are filled with a variety of large dense-core vesicles of the type usually storing and releasing neuropeptides and non-classical neurotransmitters (Zhu et al. 1986; Leng and Ludwig 2008; van den Pol 2012; Witvliet et al. 2021). A variety of neuropeptides and hormones have been identified in the *Octopus vulgaris* VL - serotonin, dopamine, octopamine, FMRFamide-related peptides, vasopressin, oxytocin, octopressin, Y-peptide and more (Shomrat et al. 2010; Shigeno and Ragsdale 2015; Winters et al. 2020; Stern-Mentch et al. 2022). Tyrosine hydroxylase (TH) - labeled processes show a wide distribution in the VL neuropil similar to that of the NM processes, suggestive of a neuromodulatory catecholamine innervation (Stern-Mentch et al. 2022).

As schematized in **Figure 10B**, the AF fibers form close associations with the SAMs (**Figure 9A**), while the CIN fibers appear to exclusively innervate the CAMs (**Figure 8, schematized in Figure 10A**). The AF boutons form a rosette-like structure in which several SAMs make adjacent synapses onto one AF bouton, mostly filled with dense-core vesicles. Such a connectivity motif between neurons and neuromodulatory processes suggests that co-activated SAMs may activate an AF bouton that, in turn, releases a neuromodulator. This mechanism can function locally, especially if the AF processes are inexcitable. Local release of the neuromodulator can heterosynaptically induce synaptic plasticity, similar to the well-known heterosynaptic serotonin-dependent synaptic plasticity, particularly in the gastropods mollusc *Aplysia* (Bailey et al. 2000; Kandel et al. 2014). The reciprocal SAM AF connection in the VL network is a plausible synaptic motif that may suggest a “self” LTP-induction mechanism as shown schematically in Figure 10B.

The feasibility of such a mechanism is hinted at by 5-HT causing both short-term synaptic facilitation of the input to the AMs and reinforcing LTP induction (Shomrat et al. 2010). Dopamine and octopamine also affect synaptic physiology but, in contrast to 5-HT, they suppress the transition to LTP maintenance (unpublished results). Immunohistochemical studies have also revealed a rich distribution of serotonin-(Shomrat et al. 2010; Shigeno and Ragsdale 2015) and catecholaminergic-rich fibers (Stern-Mentch et al. 2022) that resembles the AF distribution described here, suggesting possible AF involvement in modulation of the activity-dependent LTP. While Shomrat et al. (2010) found serotonergic immunoreactive cell bodies outside the VL and in the subvertical lobe, Stern-Mentch et al. (2022) described a group of TH-labeled cell bodies organized as a “deep nucleus” at the border between the SFL and VL. This suggests that, while incoming serotonergic innervation may provide supervision signals, like reward and punishment (Turchetti-Maia et al. 2017), a dopaminergic system may be involved in the regulation of the internal state of the SFL-VL system.

The exclusive innervation of the very small group of CAMs by a distinct group of processes (CIN) must be functionally significant (schematically shown in **Figure 10A**). The CIN processes resemble fibers that Young speculated were ascending fibers carrying “pain signals” from the arms and mantle (Gray, 1970; Gray and Young, 1964). Unlike the NMs and AFs that barely display any presynaptic profiles, the CIN outputs to the CAMs have the typical morphological features of fast transmitter synapses (**Figure 8**). The presynaptic features are similar to vertebrate presynaptic terminals, with small electron-dense synaptic vesicles, which are occasionally associated with inhibitory synapses (McDonald et al. 2002; Kasthuri et al. 2015). We therefore speculate that, in addition to feedforward excitation from the SFLs and SAMs, the CAMs integrate inhibitory inputs from the CIN. The CINs may provide negative feedback adjusting the level of LNs output to that of its targets, such as the subvertical lobe to where the LNs axons project and which Young described as the origin of the “pain fibers.” In the framework of the VL model (Shomrat et al. 2015; Turchetti-Maia et al. 2017), in which the GABAergic LN output (Stern-Mentch et al. 2022) inhibits the octopus’ tendency to attack, the CIN inhibitory input onto the CAMs would increase the LN output, resulting in suppression of the attack behavior. We believe this is what occurs when training octopuses on a passive avoidance task (Shomrat et al. 2008).

### Many SAMs together with a few CAMs appear to be organized in a canonical “association module”

Our reconstructions highlighted that ensembles of closely intermingled groups of SAM and CAM neurites form a *quasi*-columnar structural unit (**Figures 7A,7C,9A,9E, Figure - figure supplement 1**). As suggested above, this group of closely packed thin neurites likely functions as an “association module”. The neurons in each module may respond to a combinatorial ensemble of sensory features from the SFL, as schematically represented in **Figure 11 inset**. Such combinatorial coding is a common feature in association networks such as the insect and mammalian olfactory systems (Malnic et al. 1999; Lledo et al. 2005; Galizia and Szyszka 2008; Turner et al. 2008; Gruntman and Turner 2013; Kurian et al. 2021). However, the one-to-one SFL-to-SAM connections suggest that each SFL neuron is represented in probably less than 12 ± 3.6 SD SAMs, most of them in different modules spread randomly over the VL cortex (**Figure 11**). This suggests that each of VL association module associates many (i.e., several hundreds of SMAs) random combinations of visual features that are represented in the 1.8 million SFL neurons. As the number of association modules in the VL is very large (tens of thousands) it may enable the VL to “learn” the huge amount of data associated with storing the visual environment. In principle the “association module” is equivalent to that of a single interneuron in the input layer of a classical “fan-out fan-in” classification network as proposed in Shomrat et al., (2011). However, the processing power provided by the association module is computationally more elaborated than a single neuron can provide. It is tempting to speculate that what is achieved in other systems by a single complex neuron is achieved in the octopus by the construction of a microcircuitry composed of many neurons with simple properties (e.g., a linear input/output relationship).

**Figure 11.**
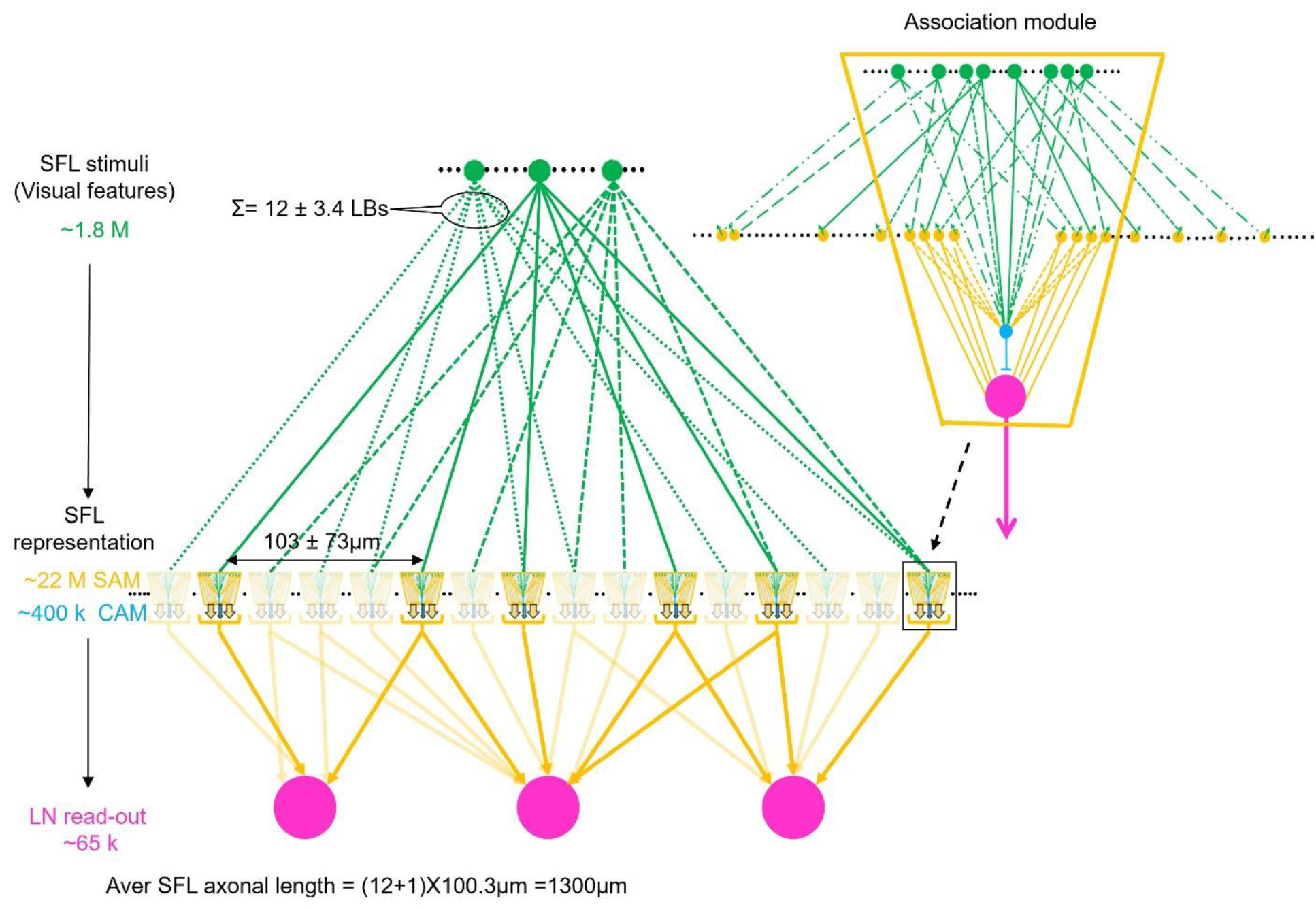
Schema of combinatorial coding in possible association modules array. *Inset:* The SAMs and CAMs appear to be organized in columnar structures that integrate and feedforward “sharpened” stimulus-specific inputs into the LNs. A specific stimulus (e.g., a visual feature) is fed forward via the SAMs while the CAMs integrate these and the SAM inputs together and feed a general balancing inhibitory input forward to the LNs. As the input to the SAMs is endowed with LTP, we suggest that this microcircuit unit is a canonical “association module” of the VL. ***The main scheme*** shows how an array of association modules that integrates a very sparse and randomly distribution of the SFL single input to the SAMs (1.0 3 ± 0.73 LBs per 100μm). Each association module contains likely several hundred SAMs and relatively only few CAMs (1.8%) suggesting that each module associates a variable combinatorial association of multiple SFL features. Note that computationally an association module replaces a single AM in the input layer of the originally assumed fan-out fan-in classification/association network (Shomrat et al 2011).

In summary, while earlier reports (Gray and Young 1964; Gray 1970; Young 1971) suggested that the VL is organized in a simple fan-out fan-in feedforward architecture much like an associative “perceptron” network (Shomrat et al. 2011), our study has revealed that its organization is more complex. Indeed, the VL circuitry is characterized by two parallel and interconnected feedforward networks that are assembled in what appear to be association modules, in which the SFL axons that convey processed sensory inputs fan-out into many SAMs and fan-in into a low number of CAMs. Both SAMs and CAMs feed the excitatory and inhibitory inputs, respectively, forward into the LN output layer.

## AKNOWLEDGMENTS

This work was supported by Human Frontier Science Program (HFSP) grant no. RGP0042/2019-102, Israel Science Foundation (ISF) grant no. 1928/15, National Institutes of Health (NIH) grant no. U19 NS104653, U24 NS109102-01, NIH 1U01NS108637. We thank Prof. Tamar Flash for her contribution during the inception phase of the project. We thank Prof. Jenny Kien for editing the manuscript.

## Author contributions

Sample preparation, F.B., Y.M., X.L.; sectioning, R.L.S.; imaging, F.B and R.L.S.; alignment, A.P. and Y. W.; manual segmentation, F.B, F.Y., X.L. and A.S.; machine learning-aided segmentation tool development, Y.M. and E.C.P.; Figures, F.B; Videos, Y.M and D.R.B.; statistical analyses., F.B. and Y.M..; 3D renderings, F.B. and D.R.B.; electrophysiological data., T.S. and B.H.; writing, F.B, Y.M and B.H.; review, D.R.B.; funding acquisition, J.W.L and B.H.; guidance, J.W.L

## Methods

### Tissue preparation

A mature wild *Octopus vulgaris* was collected by local fisherman on the Israeli Mediterranean coast and was held in artificial seawater in an individual aquarium for a few days. As previously published (Shomrat et al. 2011), the animal was deeply anesthetized in seawater containing 55 mM MgCl2 and 2% ethanol. The supraoesophageal mass was removed through a dorsal opening of the cranium (Young 1971) and then quickly sliced into 1 mm sagittal sections using a mouse matrix brain slicer and double-edged razor blade. Three slices containing the vertical lobes were obtained and immersed in fixative solution (20:1 ratio).

### Fixation

Because the preservation of the ultrastructure is a central requirement of a connectomics study, 3 fixations protocol were tested. The 3 solutions were chosen for their various aldehyde concentrations and tonicity as well as their previous use with octopus or other marine invertebrate tissue preparations for electron microscopy. Each slice was immersed in one of the following:

<u>Fixative #1</u> (Kier 1996): 3 % glutaraldehyde, 0.065 M phosphate buffer(PB), 0.5% tannic acid and 6% sucrose for 12 hr at 4°C.
<u>Fixative #2</u> (Feinstein et al. 2011): a mixture of 3 % glutaraldehyde and 4 % paraformaldehyde formaldehyde in artificial seawater for 4 hr at room temperature (RT).
<u>Fixative #3</u> (Schmidt and Derby 2011): 5% glutaraldehyde in 0.1 M PB containing 15% sucrose for 4 hr at RT.

After fixation, the slices were rinsed in 0.1M PB 4 × 30 minutes, then transferred to chilled 0.065 M phosphate buffer with 1% glutaraldehyde and 6% sucrose and stored at 4°.

We chose fixative #1 for the volumetric reconstruction, constituting the ‘large connectomics dataset’ (4nm/px, 200 ns/px), due to good preservation of the macroelements *(i.e.*, cell body) and ultrastructure *(e.g.*, mitochondria, postsynaptic density, various types of synaptic vesicles, etc.) as well as the high contrast between elements. Fixative #3 showed poor preservation of the macroelements but greater expansion (more extracellular space) and overall better fine ultrastuctural preservation. Accordingly, this fixative was used to generate a small ‘high-magnification dataset’ (1nm/px, 2 μs/px) to examine possible synaptic specialization and the synaptic vesicles at the sites of LTP generation in the VL (Hochner et al., 2003, Shomrat et al., 2011).

### EM staining

The samples were stained with a modified *en bloc* staining protocol that was developed by Hua et al. (2015) for volumetric EM imaging. Before staining, the chemically fixed tissues were rinsed 3 times for 20 min each in 150mM sodium cacodylate buffer (pH 7.4). The tissues were then stained in 2% w/v

OsO4 buffered with 150mM cacodylate for 2 hours. After being rinsed 3 x 20 min in cacodylate buffer, the tissues were immersed in 1.5% w/v potassium ferrocyanide buffered with cacodylate for 4 hours. The tissues were rinsed in deionized water for 3 x 20 min and then incubated in filtered 0.5% w/v thiocarbohydrazide aqueous solution for 1.5 hours. The tissues were then rinsed in deionized water for 3 x 20 min and stained again with 2% w/v OsO4 aqueous solution for 3 hours. After the second osmication, the tissues were rinsed in water for 3 x 20 min and transferred to 1% w/v uranyl acetate aqueous solution which was wrapped in aluminum foil to protect from light for overnight staining (approx.12 hours, RT). On the next day, the samples had a final rinse in water and were then dehydrated through 25%, 50%, 75%, 100%, 100% acetonitrile solution for 20 min each into pure acetonitrile. The dehydrated samples were infiltrated with 25% EPON:acetonitrile for 6 hours, 50% EPON:acetonitrile for 12 hours, 75% EPON:acetonitrile for 12 hours, 100% EPON for 12 hours, 100% EPON for another 12 hours. The fully infiltrated samples were placed in a flat embedding mold and incubated in a 60°C oven for two days.

### Collection of serial ultrathin sections using an automated tape-collecting ultramicrotome (ATUM)

The block face of the sample cylinder was trimmed into a 1 × 1 mm square containing the vertical lobe. 30 nm thin sections for the ‘connectomics dataset’ and 40nm sections for the ‘high magnification dataset’ were cut using a Leica ultramicrotome and collected on a home-built automated tape collecting system using a plasma-treated polyamide 8 mm wide tape (Kapton, Sheldahl) that was carbon-coated (Hayworth et al. 2014). On collection of the sections, the tape was cut into strips, and these were mounted on 100 mm silicon wafers. The connectomics dataset was composed of 7 wafers, between 115-140 sections per wafer, (total 892 sections), while the high magnification dataset was composed of a single wafer of 83 sections.

### EM imaging

For the connectomics dataset the sections were first mapped at low resolution (**Figure 2A)** and the EM image stack was aligned by Fiji’s plugin “Linear Stack Alignment with SIFT” to form a coherent 3D EM volume. The VL region free from large cracks in the tissue was chosen for high resolution imaging. The sections on the wafer were cleaned while in the Magellan microscope using the FEI-supplied plasma system. A FEI Magellan Scanning Electron Microscope (Thermofisher Scientific) equipped with custom image acquisition software (WaferMapper, Hayworth, Morgan et al. 2014) was used for high resolution imaging. A backscattered electron detector in long working distance mode with a ~3.2 NA current at 7 kev incident electron energy and a dwell time of 200 ns/px was used for imaging at 4 nm/px. For each section we obtained a 4 by 6 montage of image tiles (each tile was a 16000 x 16000-pixel image). At the end of each imaging run, out of focus images were manually identified and retaken. Artefacts such as dirt, lead precipitates, wrinkles or even large folds were seen on many sections and severely damaged sections were removed from the total stack. This resulted in 51 sections missing from the 892 sections. However, the occurrence of serious damage in adjacent sections was rare, allowing confident tracing of the elements.

### Image stitching and alignment

The stitching and alignment of the dataset were performed using a method inspired by Saalfeld et al (2012). Each step was parallelized over image tiles and sections. The single-beam electron microscope produced high resolution image tiles sized 16,000×16,000 pixels whose precise location relative to one another was not known at the nanometer level. Hence, stitching these tiles together required using image information. We stitched these image tiles for each of the 892 sections. The process consisted of a coarse-grained stitching followed by fine-grained stitching. The coarse-grained stitching performed a per-tile affine transformation, setting the tiles roughly in their final position. SIFT features (Lowe 1999) were computed for pixels up to 2,400 from the tile boundary. The features of the overlapping tiles were compared and matched to find point correspondences. To remove outliers of incorrect correspondences, the RANSAC algorithm (Fischler and Bolles 1981) was used, optimizing for rigid transformations. Finally, the tiles’ affine transformation was optimized by minimizing the sum of squared roots of all correspondences.

To achieve better alignment and compensate for minor non-linear deformations in imaging, a fine-grained stitching was performed after the coarse-grained stitching. A triangular mesh was laid on each tile and a 1×1 μm2 area around each mesh vertex overlapping with an adjacent tile was cropped. The cropped patch was matched against a transformed cropped area of the neighboring tile of size 2×2 μm2. The 2D elastic optimization algorithm described in Saalfeld et al. (2012) was used to minimize the distances of the matches.

We applied a custom-made 3D alignment algorithm to obtain an image stack of all 892 sections. Once all the sections were stitched as described above, correspondences between adjacent sections were detected. The first step consisted of coarse features detection and matching, and subsequently patch matching in small regions. The matching was performed between all section pairs separated by at most one section (hence considering matches between non-adjacent sections). Coarse features were obtained from the stitched sections. The features were detected using the OpenCV SimpleBlobDetector (“OpenCV: cv:SimpleBlobDetector Class Reference” n.d.). The detector searched for circular blobs that typically corresponded to blood cells, mitochondria and axonal cross-sections. Features were defined using a SIFT descriptor (Lowe 1999). After matching the features of each pair of sections, RANSAC filtered out wrong matches, from which an affine transformation between the sections was computed. In the second step, finegrained patch matching at half resolution was performed. A triangular mesh was laid on the sections and a 1.6×1.6 μm^2^ area around each mesh vertex was cropped, followed by cross-correlation search against a transformed cropped area of the neighboring section of size 4×4 μm2. The valid cross-correlation matches were then used as an input to a 3D elastic optimization algorithm based on Saalfeld et al. (2012) which minimizes the cross-layer distance of the matches while preserving the 2D structure of the sections.

### Semi-automatic segmentation: mEMbrain

To segment the aligned image stack (892 sections, volume 2.7 million μm^3^; **Figure 1B**), we applied an automated reconstruction pipeline to the volume using a machine learning reconstruction pipeline (Lee et al. 2017; Meirovitch et al. 2019). The inputs to this pipeline required fully annotating the image stacks of 4 manually picked volumes (image stacks ranged in volume from 21.6 to 153 μm^3^; **Figure 1-supplement 3**), which served as ground truth for training the artificial neural network. The annotation of this ground truth labeled two different categories: intracellular space, excluding cellular membranes (category 1), or any combination of cellular membrane and ECS (category 2). We leveraged the mEMbrain software package implemented in MATLAB (Pavarino et al. 2019) to obtain a first template of the cellular processes. This computation included training a deep neural network (Unet; Ronneberger et al., 2015) to classify pixels according to the two categories (accuracy above 93%) and produce a 2-dimensional instance segmentation of the individual cross-sections of all cellular compartments using Watersheds (Pavarino et al. 2019). We used custom code to merge colored neuronal cross-sections across slices of the image stack if adjacent cross-sections matched in shape or if adjacent processes had sufficient overlapping cytoplasm far from any cellular membrane. Applying this conservative merging resulted in a 3-dimensional representation of object instances that each rarely overlapped two distinct neuronal or glial processes, and often the splitting of complete neuronal processes into several segmented objects. In most cases, however, the 2-D cross sections were lawfully (fully) segmented into one object and after conservative agglomeration procedure based on shape matching these objects were correctly agglomerated over a few consecutive sections. We also applied 3D agglomeration procedures including mean affinity agglomeration suggested by Lee et al. (2017) and Meirovitch et al. (2019). However, due to membrane breaks these procedures often resulted in merge errors that were hard to manually proofread and hence this agglomeration was not included in the circuit analysis. We found the cause of the merge errors to be associated with specific cellular and compartment types, including the shafts of the large neurons and many synaptic boutons. Because such preparation artifacts severely affected the quality of our fully automated reconstruction, we combined the automated segmentation with manual methods described below.

### Membrane-constrained ML-aided reconstruction

To facilitate manual reconstruction we used the mEMbrain package (Pavarino et al. 2019) implemented in MATLAB (Mathworks) together with VAST (Volume Annotation and Segmentation Tool (https://software.rc.fas.harvard.edu/lichtman/vast/, Berger et al., 2018). Using U-net architectures of deep neural networks (Ronneberger et al. 2015) mEMbrain computed the boundaries of all cellular processes, scoring the likelihood (0 to 1) that pixels belonged to intracellular space (category 1) or not (category 2). mEMbrain arranged these scores into a volume of membrane probabilities consistent with the EM image stack layer in VAST, which allowed tracing cellular processes aided by the pre-computed membrane probabilities. We constrained the tracing to the membrane probabilities layer generated by mEMbrain using the “constrained painting” function in VAST, which restricted the tracing to connected components of pixels far from cellular membranes. This ML-aided tracing allowed the annotator to ignore the location of the cellular membranes and work in a skeletonization mode if the initial painting stroke overlapped intracellular space. Tests on small volumes suggested a ~5-10x faster annotation in this mode than manual painting without constraints, while retraining the accuracy of careful pixel annotation. This membrane-constrained ML-aided reconstruction was used for the 516 cells described and studied morphologically here (**Figure 1C**).

### Node-based agglomeration of 3D compartments

We also converted the automated neuron reconstruction IDs into a VAST image stack layer holding the identities of ~16×10^6^ objects to allow even faster semi-automated reconstruction of processes. In this procedure the annotator used the automatic reconstruction by iteratively agglomerating 3D objects that were found to belong to a single cellular process. This was done using VAST’s “Tool Layer” functionality which we mapped to the automated segmentation layer and to an additional annotation layer that represented the annotator’s manually made skeletons. The annotator placed single nodes within the objects segmented by the automated reconstruction pipeline; together the collection of nodes of a single skeleton defined the agglomeration of all objects straddled by the nodes into a single proofread cell. This membrane-constrained ML-aided reconstruction was used to quickly reconstruct all the synaptic partners of one LN (**Figure 7C**) and one AF (**Figure 9 - figure supplement 1**).

### Criteria for cell classification

Our morphological reconstruction included 516 cells segments and a wiring length of more than 12.8 cm. Only neurons that were sufficiently contained within the reconstructed volume were classified. Processes that could not be followed across the volume or were not wholly contained within it were considered orphan fragments and fell into the ‘unmarked category’ with other uncharacterized profiles.

#### 1. Afferent axons

Classified as afferent axons were processes of constant radius (as opposed to dendrites that often taper), with exclusively presynaptic boutons filled with synaptic vesicles smaller than 60 nm and with no connection to a cell body. Two processes met these features: the SFL axons (see Results; **Figure 3** and the CIN (see Results; **Figure 8**). The SFL axons were classified according to several established criteria (Gray and Young 1964; Gray 1970; Young 1971): the axon projected into the VL lobule within the outer zone of the neuropil, made *en passant* synapses with AM neurons and at the ultrastructural level their varicosities showed an electron-lucent cytoplasm with average synaptic vesicle size of 59.7 ± 1.5nm, and numerous round mitochondria. Unlike the SFL, CIN were found in the SFL and in the neuropil, displayed the smallest vesicles in our dataset (38.2 ± 0.9 nm) and synapsed exclusively with clear postsynaptic profiles. Additionally, CIN fibers showed a high concentration of neurofilaments which ended where the synaptic vesicles appeared (Gray 1970). The high neurofilament content of these processes led Gray (1970) to suggest that these were pain fibers. Both afferents could be easily identified with only electron micrographs.

#### 2. VL Interneurons

Neurons were classified as VL interneuron if they received input from the SFL and/or were previously described as intrinsic to the lobes. All partners of the SFL axons were the AM cells (Gray and Young 1964; Gray 1970; Young 1971). However, we discovered two types of AMs: SAM and CAM (see Results; **Figure 6**). The SAMs showed a single inwardly directed trunk that cannot be defined either as axon or dendrite. Additionally, their ultrastructural signature was easily identifiable through the presence of synaptic vesicles (49.5 ± 1.3 nm) along the length of this trunk and the presence of specific ultrastructural components such as cisternae and the transport endoplasmic reticulum at every presynaptic site. Unlike the SAM, the CAM displayed a strict segregation between a dendritic compartment located within the SFL tract and their axonal process within the neuropil. Their dendritic tree was lightly or heavily branched, with one or multiple axons arising from it. Their dendritic arborizations displayed a classical electron-lucent postsynaptic profile filled with polyribosome. Unlike the SAMs, the CAM presynaptic sites in the neuropil displayed a clearer cytoplasm with no specific ultrastructure and smaller vesicles than the SAMs (42.6 ± 1.1 nm).

We also observed putative interneurons displaying a bifunctional neurite, with both pre- and postsynaptic sites identified (**Figure 1-figure supplement 3A**). Previously undescribed and poorly represented in our dataset and apparently poorly involved in the major cell network, all such cells were considered as unmarked cells in the data analysis.

#### 3. Efferent Fibers

Only the LNs fell into this category. Their cell bodies (diameter > 15 μm; n=14) lay mostly in the inner zones of the cortex and their large trunks ran radially into the neuropil (see Results; **Figure 7**). While most of the LNs were poorly contained in the reconstructed volume, some gave off dendritic collaterals within the SFL tract area where they were solely postsynaptic to the SAMs and unmarked process. Their dendrites showed a variable diameter along their length and displayed a typical clear postsynaptic profile. In the neuropil, we classified all clear postsynaptic profiles with convoluted arborizations heavily innervated by the AMs as LN processes.

#### 4. Neuromodulatory neurons

Based on known invertebrate modulatory synapses, processes were classified as modulatory if they contained mostly large vesicles (>60 nm), either dark or clear. We distinguished two types of neuromodulatory processes (see Results; **Figure 9**). First, the ascending fibers (AF) with a distinctive morphology - a menorah –shaped tree climbing from the neuropil to the cortex while intermingling with SAM trunks. Along their length the AF displayed boutons filled either with dense core vesicles (AF-Nm) and an average size of 74.9 ± 1,6 mm or a mix of smaller vesicle types that could be round, flattened or irregularly shaped (AF-mix). Various subtypes of AF-mix were observed. The AF-Nm neurons were heavily innervated by the SAMs without any sign of presynaptic activity. In contrast, the AF-mix were innervated to lesser extend by the SAMs and displayed some presynaptic sites as well as occasionally reciprocal contacts.

The second type were neuromodulatory fibers (NMs) which were dispersed more globally in the VL neuropil and within the SFL tract and lacked a general orientation. Several subtypes were characterized by the homogeneity of the vesicle types found along their length. Most of their boutons lacked any sign of synaptic connection but the few pre- or postsynaptic sites observed were unorthodox. These NM processes displayed the largest vesicles in the entire dataset with diameters up to 200 nm (mean 96.3 ± 3.5 nm) within their bouton and an accumulation of much smaller vesicles was observed at their rather rare presynaptic sites.

#### 5. Glia

Cells were classified as glia if they showed clearly distinguishable gliofibrils (Gray, 1964) and/or did not receive any synapses. At least two types of protoplasmic glia were identified (**Figure 1-figure supplement 3C**). (1) radial glia (Young, 1971) were full of gliofibrils and characterized by long radial processes and arborizations infiltrating the neuronal cell cortex. These cells appear to isolate neuronal types from each other. (2) The astrocyte-like glia had many gliofibrils within their processes, which showed an increased tortuosity densely infiltrating the SFL tract. Here they appeared to wrap distinct synaptic areas. Through repeating folds this cell type appears to maximize self-contacts characterized by apposition of their membranes and accumulation of electron dense materials on both sides, possibly sites for protein exchange. One astrocyte-like cell with its cell body located near a cell island was partially reconstructed; it covered an extensive area within the SFL tract. (3) Fibrous glia with cell bodies in the inner cortex were also observed; these did not receive any synapses in the reconstructed volume.

#### 6. Progenitor-like cells

Cells were classified as progenitor-like only if their cell body was reconstructed (**Figure 1-figure supplement 3B**). These cells were nested in each other in the inner cortex forming a type of horizontal layer. There were at least two types of cells in this arrangement. First, the base was formed by elongated cell bodies whose nuclei showed bumpy irregular membranes often with nuclear folds. These cell bodies appeared to have very small projections, and, like the glia, they lacked synaptic connections. Above these elongated cells were more classically spherical cell bodies which appeared somewhat “immature.” Their trunks, extending radially into the neuropil, often lacked synaptic connection and terminated early with filopodia typical of axon growth cones. Their fibers were rich in grayish filaments that could be filamentous actin, suggesting that this may be a site of plastic reorganization. Because of the similarity of this arrangement with neurogenic niches observed in the mammalian dentate gyrus of the hippocampus and the subventricular zone, we termed these cells “progenitor-like cells.” We saw no sign of cell division.

### Identification and analysis of synapses

Chemical synapses were identified by features common to invertebrate synapses - the presence of a cluster of synaptic vesicles (sv) or ‘cloud’ associated with a lightly active zone near the presynaptic membrane (Gray 1970; White et al. 1986; Ryan et al. 2016; Zheng et al. 2018; Meinertzhagen 2019; Witvliet et al. 2021). Synapses were annotated if the accumulation of sv persisted through at least 3 sections in a row (90nm). We distinguished between traditional chemical synapses (fast transmission) with 30-60nm sv and synapses of modulatory neurons with mostly large vesicles (Witvliet et al. 2021). Consistent with previous studies (Gray and Young 1964; Gray 1970), we did not find evidence of electrical connections (gap junction) or any clear postsynaptic density (PSD) at any chemical synapse within the vertical lobe. However, in the connectomics dataset we observed a rather global membrane thickening at synaptic sites. This may be partly due to a lack of resolution and the observed global overstaining.

The high magnification dataset was used to confirm the synaptic nature of the connections observed in the connectomics dataset and the absence of PSD. The absence of PSD made it difficult to annotate polyadic varicosities (one cluster of vesicles could be associated with two postsynaptic targets). In ambiguous cases only the most likely partners were identified (larger surface contact), leading to a likely underestimation of the overall number of synapses. Because most of the cells were not fully contained in the reconstructed volume, it was not possible to assess the full collection of synapses between a given presynaptic and postsynaptic neuron. For this reason, we refer to “synaptic site,” a single anatomical release site or a single synapse. Multiple synaptic sites from the same cell were classified as a single input. Additionally, because cells had different lengths of arbor in the reconstructed volume, we expressed most of the results as synaptic fraction and number of synaptic sites per 100 μm.

Although multiple annotators would have been ideal, due to project constraints and the difficulty of training people in an entirely new dataset where most synapses are being described for the first time, all synaptic annotations were made by a single annotator, giving consistency across annotations.

### Clustering of cells based on connectivity

As described above, the reconstructed cells and processes of the vertical lobe were classified based primarily on morphological criteria. We combined this classification with unsupervised clustering to obtain a visual summary of the connectivity patterns found in the lobe. First, we produced a cell type wiring diagram connecting the 7 main cell types (**Figure 2**). The nodes of this graph comprise cell types and the weighted edges between pairs of cell types represent the number of synapses between the processes that belonged to these cell types. We used a spring model to minimize the distances between the pairs of nodes (representing pairs of cell types) that corresponded to a relatively larger number of synapses (Fruchterman and Reingold 1991). This diagram revealed that synapses within a cell type are rare in the vertical lobe (mainly in ascending fibers; see Results). In addition, visually inspecting this diagram clarified that the connectivity was highly structured and far from an “all-to-all” connectivity. Each cell type was connected to a small number of other cell types (see Results). This observation led us to consider the common inputs and common outputs of specific processes (described below).

We also discovered that besides these uncommon within-type synapses, there were no connectivity cycles in the cell-type wiring diagram. This observation led us to analyze the possible cycles in the wiring diagram while considering individual processes (see above Criteria for cell classification). We then used an unsupervised approach to visualize the connectivity among individual processes (**Figure 2 – figure supplement 1**). This was done by considering the adjacency matrix **A_NxN_** whose **(i,j)** entries represent the number of synapses from the presynaptic neuron **i** to the postsynaptic neuron **j,** considering **N** neurons (**N**=362). As specific groups of cells tended to synapse into other specific subsets of cells, a stereotypy observed from the cell-type diagram, we considered the number of shared inputs and shared outputs between individual processes, mathematically represented by the common output and common input symmetric matrices, **A*A’** (**presynaptic**) and **A’*A** (**postsynaptic**), respectively. The presynaptic matrix represents how many common targets exist for pairs of neurons and the postsynaptic matrix represents how many common inputs exist for pairs of neurons. To visualize the tendencies of neurons to share inputs or outputs, we considered the matrix **A*A’+ A’ *A** which sums for every pair of neurons the number of shared inputs and outputs, using subspace embedding (Koren 2005) and 100 dimensions, heuristically picking a dimension that yielded sufficient separation between individual nodes (nearby dimension produced visually similar outcomes). This embedding of the neurons in the plane resulted in a close positioning with other neurons with a similarity of inputs and outputs (summed). The position of the entire layout and its orientation are arbitrary and not used to derive biological observations. To combine the cell type- and neuron-based diagrams, each neuron was located based on its type diagram shifted by its location in the neuron-based wiring diagram (**Figure 2**).

### Analysis of feed-forwardness

The cell type-based wiring diagram analysis revealed that connectivity cycles were rare and information flowed in one direction among populations of cells belonging to different cell types (**Figure 2**). To determine if there were cyclic synaptic information flows in the reconstructed circuit at the level of individual neurites we used graph algorithms that detect directed cycles in directed graphs. Topological sorting algorithms (Barth et al. 2004) that determine a linear order in cycle-free directed graphs were used to detect cycles in the graph. A topological sorting is a sequence of nodes each of which is possibly connected only to nodes proceeding in the list. Hence, we inspected all the edges in the topological sorting that went backward. We found a few edges (n= 4) that violated the forwardness of the circuit and each of these cases was related to a reciprocal synapse between a pair of nodes associated with a single synaptic site between the cells. We also calculated the probability of observing this level of forwardness in random graphs that agreed with the wiring diagram, but in which the direction of synaptic transmission was randomized. This showed that the level of forwardness in the data is unlikely to occur by chance (p<0.05).

### Classification of cell types from cell body morphology

Our reconstruction revealed that the SAM, CAM, and LN cell bodies lay in the VL cortex along with several other cell types: glia, progenitor-like and mix cells (Others; **Figure 1 – supplement 4**). We noticed, consistently with early studies, that the LNs had extremely large cell bodies and that the newly discovered CAMs tended to have a slightly larger cell bodies than the SAMs. We tested whether these observations could be used to classify cell types directly from the morphology of their cell bodies. We used 110 cells whose somata were contained in the volume and their processes were distinctly characterized in the neuropil by an expert annotator: 7 LNs (6.36%), 69 SAMs (62.73%), 16 CAMs (14.55%) and 18 others (16.36%). The selection of this subset of cells was agnostic to the details of the classification process described here. For each of these cells we considered the volume of the cell body as the number of annotated voxels multiplied by the volume unit of a voxel, and the volume, surface area and sphericity (Wadell 1935) of the ellipsoid best matching the geometry of the cell body. Ellipsoid approximations were computed from the eigenvectors and eigenvalues of the covariance matrix of the voxel coordinates of the cell bodies.

Our tests showed that the information gained from these ellipsoids were sufficiently discriminative and hence these 3 parameters (volumes, surface area and sphericity) were used for further analyses. Support Vector Machines (SVMs) were used to train, cross-validate and predict the 3 main cell classes found in the cortex: LN, SAM and CAM and a 4th class, Other, to represent the seemingly small number of other cell types in the cortex (including glia, progenitor cells and possibly other cells of scarce representation). The min, max, mean and std of these parameters were computed for each class as well as the training accuracy (see Results).

In addition, a set of 49 cells (each cell was annotated twice) was used to determine the accuracy of the SVMs and for leave-one-out cross validation (see Results). Once the SVM model was determined from the training set, it was applied to a test set, comprising 1447 cells, that were densely annotated in the cortex (using mEMbrain and VAST as described above). The number and percentage of each of the 4 classes were calculated (see Results; **Figure 4**).

### Wire-length calculation

To calculate the length of the processes (see Results; **Figure 2**), we used a thinning algorithm to map each volumetric object into a skeleton and used graph theory algorithms to measure the total length of skeletons. This was done by calculating the graph distance between skeleton nodes representing the end of the volumetric object (a leaf; a node that is connected to at most one other node, e.g., of the last node in a branch terminating at a synapse) and skeleton nodes representing branching (nodes that are connected to more than two other nodes, e.g., a branching point of a dendrite-like structure or axonal branching). The distances between adjacent skeletal nodes were defined using the Euclidean distance between the voxels in the 3D 26-neigberhood mask (a 3×3 cube around each voxel). We then summed up all these distances to measure the wire-length of each process. Occasionally the thinning algorithm produces a biologically meaningless cycle due to topological holes in the annotated objects (either biological or imperfect annotation). We guaranteed that only the shortest paths between special skeletal nodes (bifurcations and leaves) were used for the wire-length calculation using minimum distance graph algorithms between leaf and bifurcation nodes.

### SFL bouton volume calculation

Synaptic boutons are morphologically easy to identify, and they were accordingly individually segmented to allow volumetric measurements (see Results; **Figure 3**). Volumes of boutons were estimated from the voxel counts of each bouton. A *post hoc* exact permutation test for independent samples compared the volumes of the SFL SB and SFL LB. The p-values were adjusted according to the sequential Bonferroni correction as described by Holm (1979).

### Subcellular reconstruction

The geometry of the SFL large synaptic boutons and the postsynaptic structures they innervate were reconstructed at the subcellular level using the high magnification dataset (see Results; **Figure 5**). Neuronal elements of interest were manually segmented on VAST. The labeled images and metadata containing the labels were exported and processed externally or accessed through the VAST API. Vesicles showing a clear synaptic center in each section were counted by outlining intracellular space containing vesicles (see more information below). Mitochondria were identified by the presence of an inner membrane with visible cristae. Transport endoplasm, large electron-lucent tubular organelles, spanned the reconstructed volume and appeared to be a continuous structure present throughout the length of the AM trunk. They showed large swellings at the presynaptic site, particularly at the ‘palm’. Discontinuous thin lucent tubular organelles that sometimes appeared as dark lines were annotated as ‘cisternae’ (Cis). It is not clear whether the Cis (up to 15 observed in one palm) were part of the larger endoplasmic reticulum network or were isolated entities.

### Counting, modelling, and measuring vesicles

An annotator painted all individual vesicles of several compartments using distinct colors per compartment. Distinct colors were used to annotate the active zones. The center of mass and areas of the vesicle cross-sections were used to approximate the centers and spherical representations of individual vesicles, from which spherical 3D mesh representations were computed. Occasionally, several vesicles of the same compartment were overlapped spatially. This required us to use distance transforms and watersheds to achieve separation of vesicles. Vesicle counts were made only after they were separated in space. Due to the section thickness (30 nm), most of the vesicles were clearly appreciated in a single section, whereas only the largest vesicles (> 60nm diameter) may have had a strong signal in two adjacent sections. The annotator attempted to avoid overrepresentation of a vesicle, when possible, but since this task is not unambiguous our counts likely represent an upper bound. Extremely small vesicle-like structures were not annotated since it was assumed that these structures were likely represented in adjacent sections.

### Rendering

For rendering of all the traced images, 3D surface meshes of labeled objects were generated from VAST using VastTools (written in MATLAB) and imported into 3D Studio Max 2020 (Autodesk Inc.) to generate 3D renderings of all the traced objects.

### Data availability

The ‘connectome dataset’ is available at https://lichtman.rc.fas.harvard.edu/octopus_connectomes with 32 nm resolution. 8 nm resolution will be provided, and the segmentation files added prior publication.

