## Supplementary figures and images for "Connectomics of the *Octopus vulgaris* vertical lobe provides insight into conserved and novel principles of a memory acquisition network"

### Figure1-video supplement 2

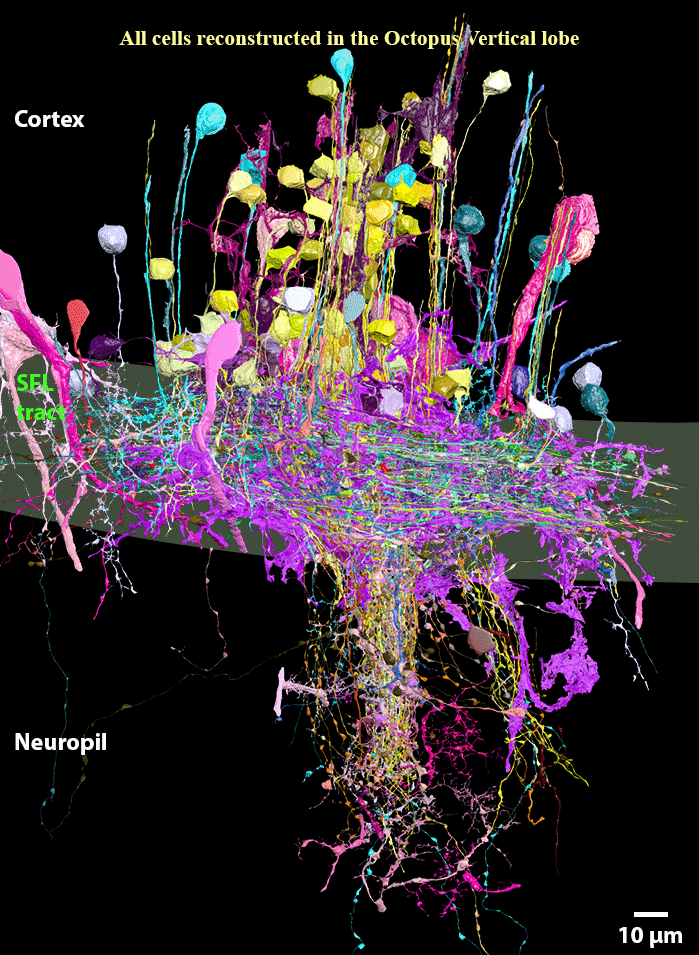
