## Supplementary Data for "Connectomics of the *Octopus vulgaris* vertical lobe provides insight into conserved and novel principles of a memory acquisition network"

**Supplementary Figure**

**Figure 1-video supplement 1.** Vertical lobe EM volume (260x390x27μm) imaged with 4 nm resolution in backscattered electrons electron mode. A ~3.2 NA current at 7 kev incident electron energy and a dwell time of 200 ns/px were used. Sections through the ROI volume are shown in sequence.

**Figure 1-video supplement 2. Vertical lobe neural elements.** Movie showing all neural elements reconstructed in the VL volume and subdivision into neural types according to morphology, ultrastructure, and synaptic connectivity. Only neurons that were sufficiently contained within the reconstructed volume were assigned a cell identity and are represented here.

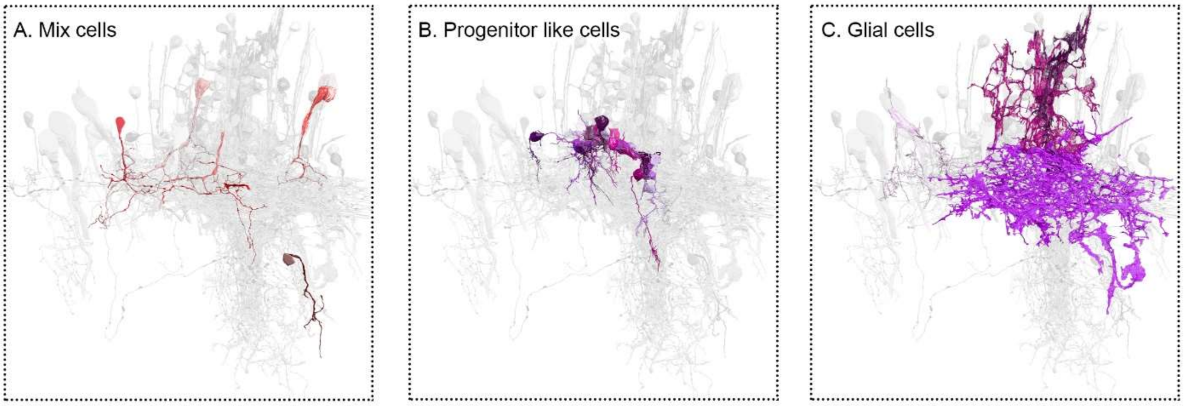


**Figure 1 - figure supplement 3. Three neural elements with yet unclear relationships with the VL connectome**

Aside from the 7 cell types highly involved in the VL connectivity our reconstruction revealed the existence of (A) putative interneurons of the VL displaying a neurite with both pre – and postsynaptic sites, which we termed 'mix cells'. They were poorly represented in the reconstructed volume and without clear involvement in the core connectivity of the VL wiring diagram. (B) In the inner cortex, cells with elongated cell body, irregular nuclear shape and a very short projection were nested in each other. Juxtaposed onto these cells were other cells with round cell body and short trunk directed inwards that lacked synaptic connections and terminated with a filopodia typical of axon growth cones. Although no cell division was observed, this organization is reminiscent of the neurogenic niche, and accordingly we termed these cells "progenitor-like" cells. (C) Although rare, we identified at least two types of protoplasmic glia cells contained within the cortex and widely spread astrocyte-like glia restricted to the SFL tract. Some putative fibrous glia were also observed.

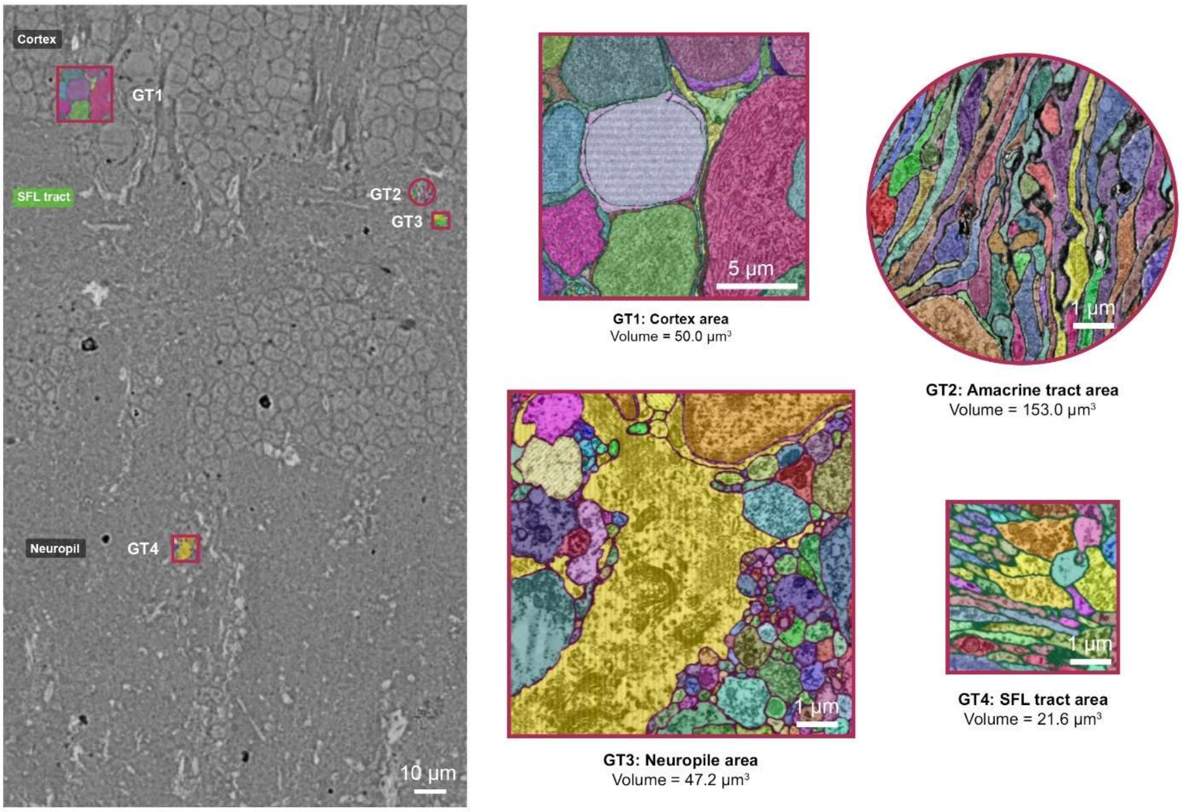


**Figure 1-figure supplement 4. Ground truth incorporated in the training dataset.** Four distinct areas were chosen to represent the diversity of the octopus VL ultrastructure: cortex, amacrine tract, SFL tract and neuropil. For each area, an expert annotator carefully painted the cytoplasm of cells using distinct colors. This procedure required careful annotation and a correct evaluation of the 3D morphology of cells. Image processing was later used to turn this multi-label set into a ground truth comprising only the boundary categories. This was used to train the deep learning model (see Methods).

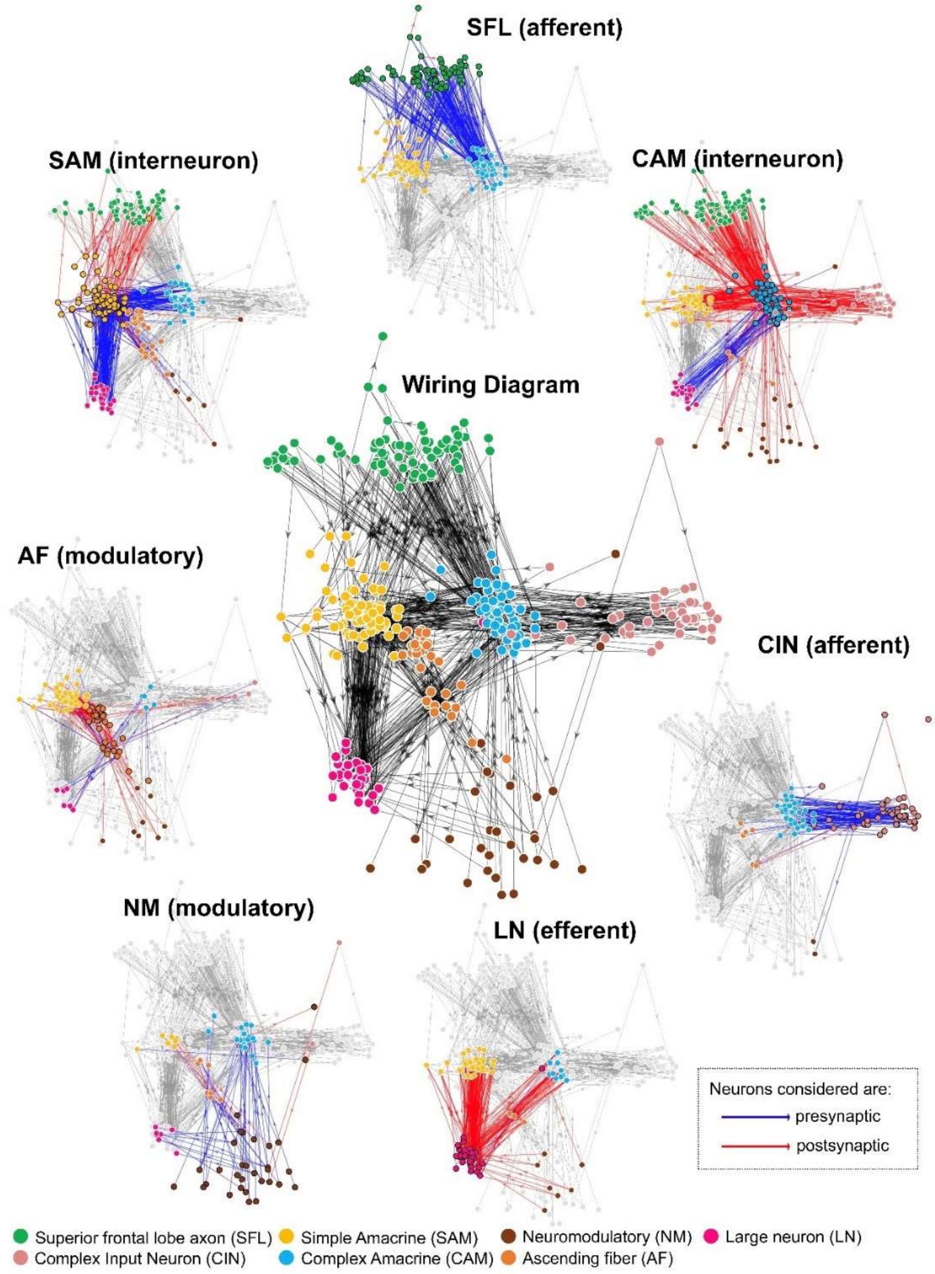


**Figure 2 – figure supplement 1. Wiring diagram at the neurite level (middle panel) surrounded by the panels of each of the 7 cell types depicting their connectivity.** Each arrow represents the direction of a chemical synapse and line color indicates the direction of connection**:** synaptic inputs (red) and outputs (blue). Connections were derived from synaptic annotations based on identified connections between the reconstructed cells and nodes were similarity positioned based on their connectivity (see Methods).

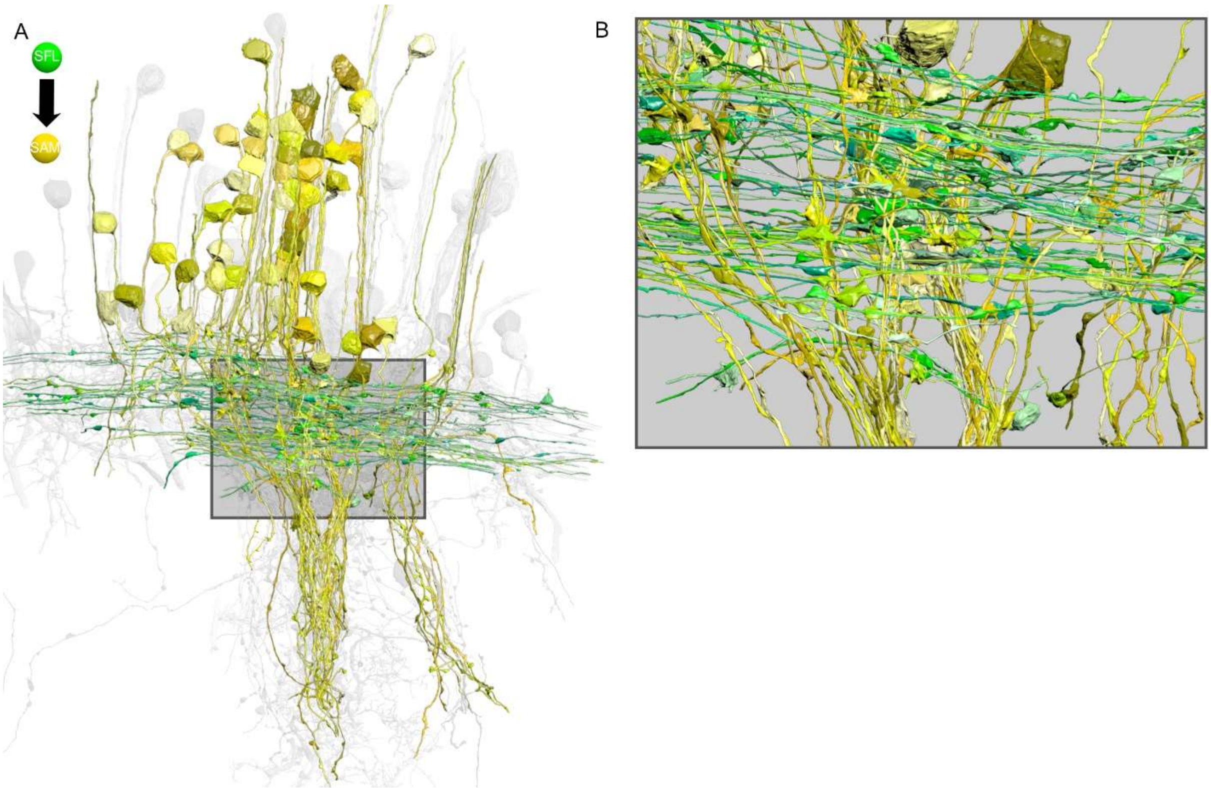


**Figure 3- figure supplement 1**. (A) The SAMs (yellow; n=108) and the SFL axons (green; n=84) form cruciform connection within the SFL tract. Notice the SFL axons running in parallel that cross the SAMs neuritic trunks perpendicularly. (B) Detail of A. The SFL large boutons can be seen synapsing onto the SAMs palm that constitute the single input of these cells. Reconstructions are superimposed on a grayscale representation of the 516 reconstructed cells.

**Figure 3-video supplement 2.** One segmented SFL axon tracked within vertical lobe EM volume (260x390x27µm) that was imaged with 4 nm resolution in backscattered electrons mode; a ~3.2 NA current at 7 kev incident electron energy and a dwell time of 200 ns/px were used.

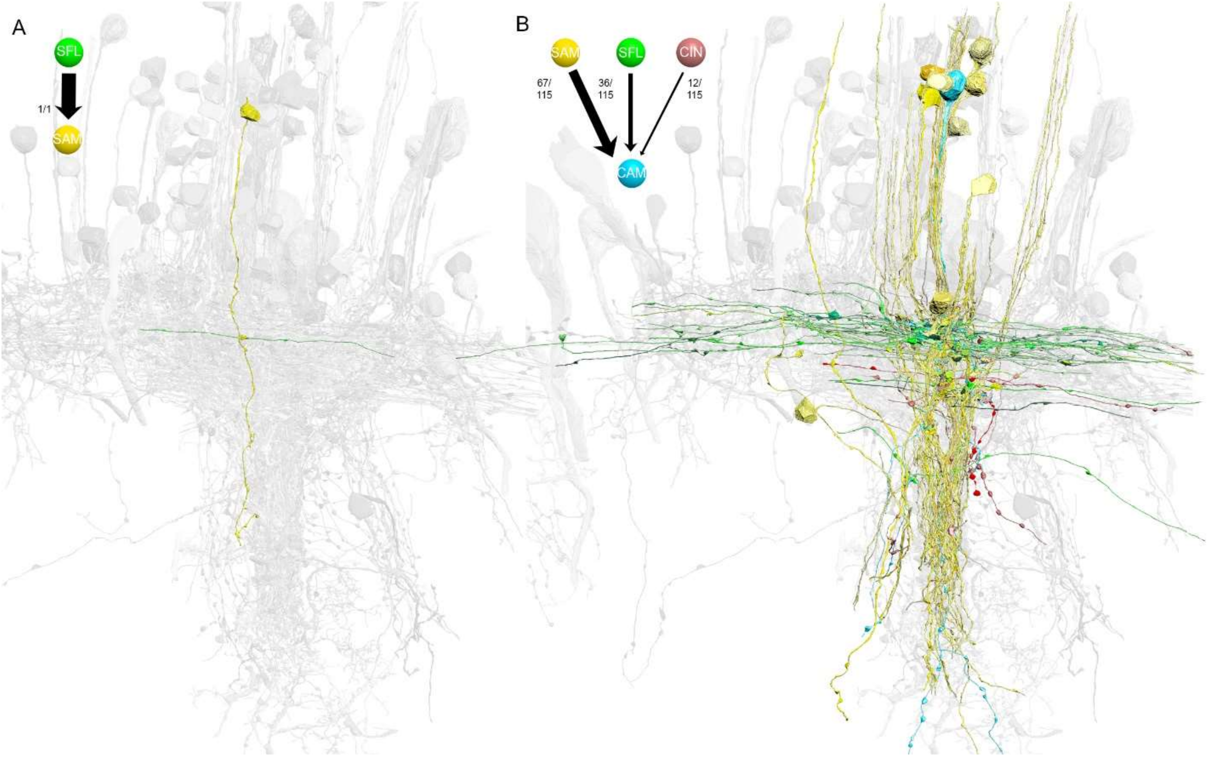


**Figure 6- figure supplement 1. Contrast between the single [mononeural] input of the SAMs and the multiple [polyneural] inputs of the CAMs.** (A) Reconstruction of one SAM (yellow) and its only input from an SFL axon (green). (B) Reconstruction of all the presynaptic partner of one CAM (cyan), 67 from SAMs, 36 from the SFL axon (green) and 12 from the CINs (violet) (total n=115). Both reconstructions are superimposed on a grayscale representation of the 516 reconstructed cells.

**Figure 6-video supplement 2.** One segmented SAM tracked within vertical lobe EM volume (260x390x27μm) imaged with 4 nm resolution in backscattered electrons mode; a ~3.2 NA current at 7 kev incident electron energy and a dwell time of 200 ns/px were used. The video progresses from the cortex passing through the SFL tract region to the neuropil.

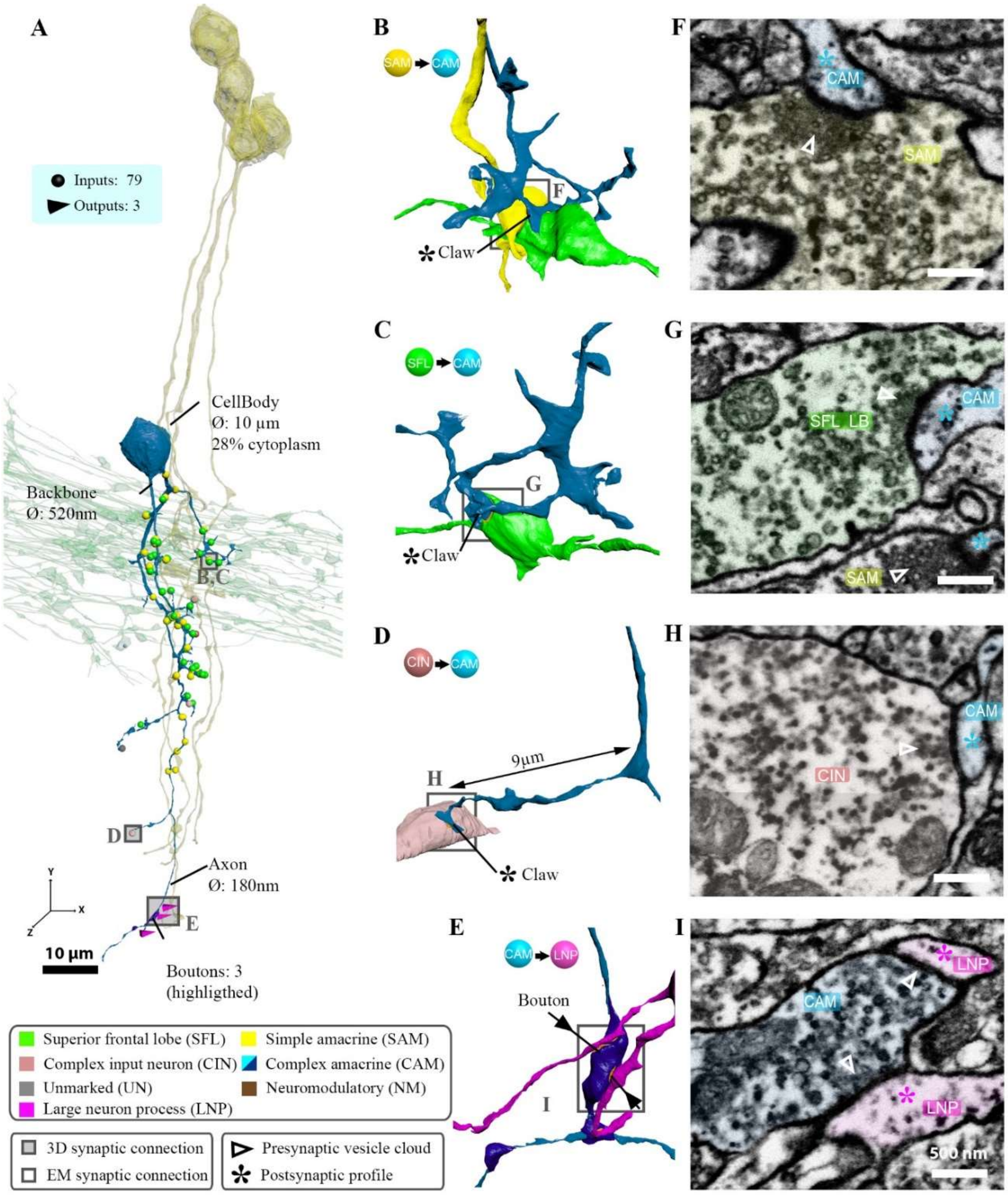


**Figure 6 - figure supplement 3. Example of a vertically projecting CAM and its connectivity.** (A) A CAM superimposed on a reconstruction of the SFL tract (green) and a few SAMs (yellow) to facilitate spatial orientation. Puncta and triangle represent, respectively, postsynaptic and presynaptic sites on the CAM, color coded according to pre- or postsynaptic partner. (B-I) Magnification insets illustrate the synaptic arrangement for the 4 main synaptic connections at the marked location together with the corresponding EM cross-section (scale bar=500nm). This CAM integrated inputs from multiple SAMs (B,F), SFL axons (C,G) and CINs (D,H) mostly within the SFL tract area. It innervated the LNp deeper in the neuropil (E,I).

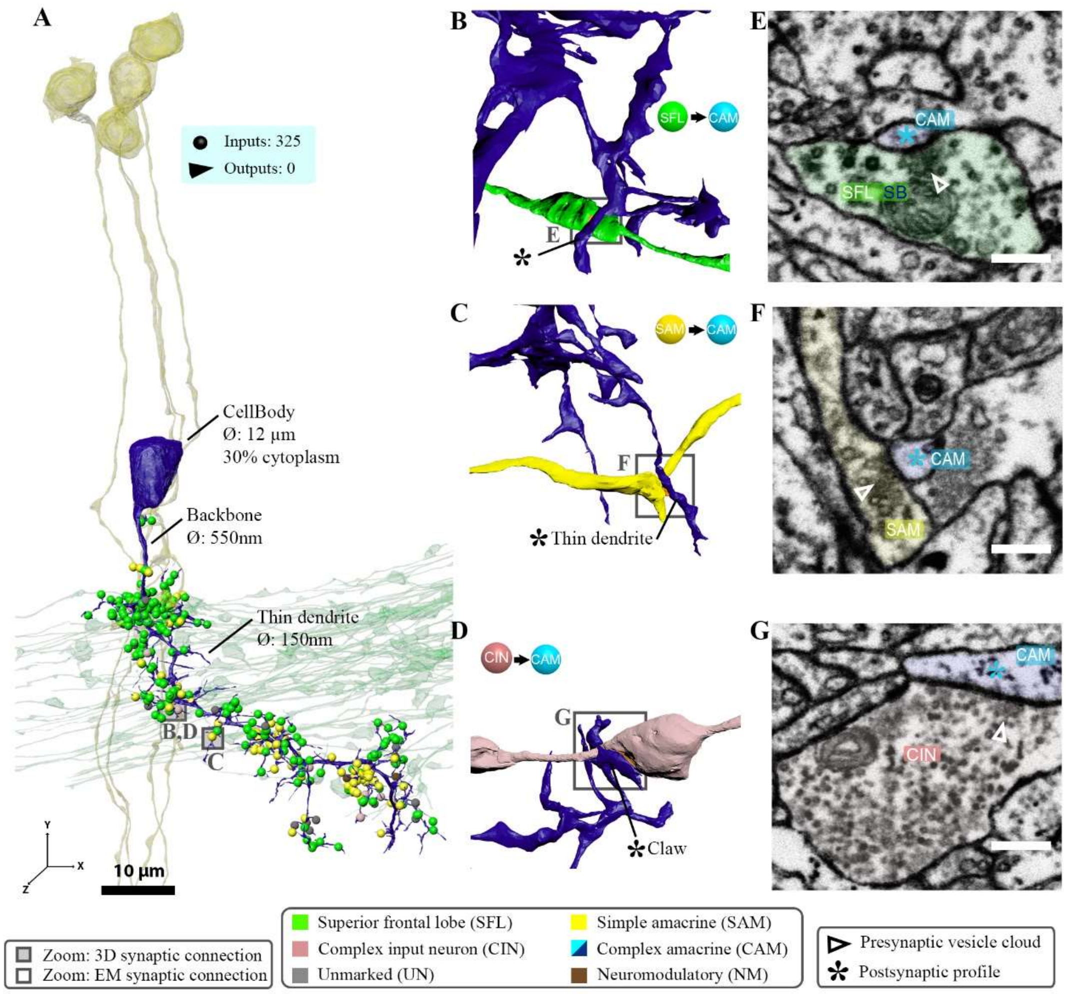


**Figure 6 - figure supplement 4. A laterally projecting CAM and its connectivity.** (A) A CAM superimposed on a reconstruction of the SFL tract (green) and a few SAMs (yellow) to facilitate spatial orientation. Puncta represent the postsynaptic input sites color coded according to the pre- or postsynaptic partner. (B-G) Magnification insets illustrate the synaptic arrangement for the three main synaptic connections at the marked locations, with corresponding EM cross-sections (scale bar=500nm). This CAM integrated inputs from multiple SAMs (B,E), SFL axons (C,F) and CINs (D,G) mostly within the SFL tract area.

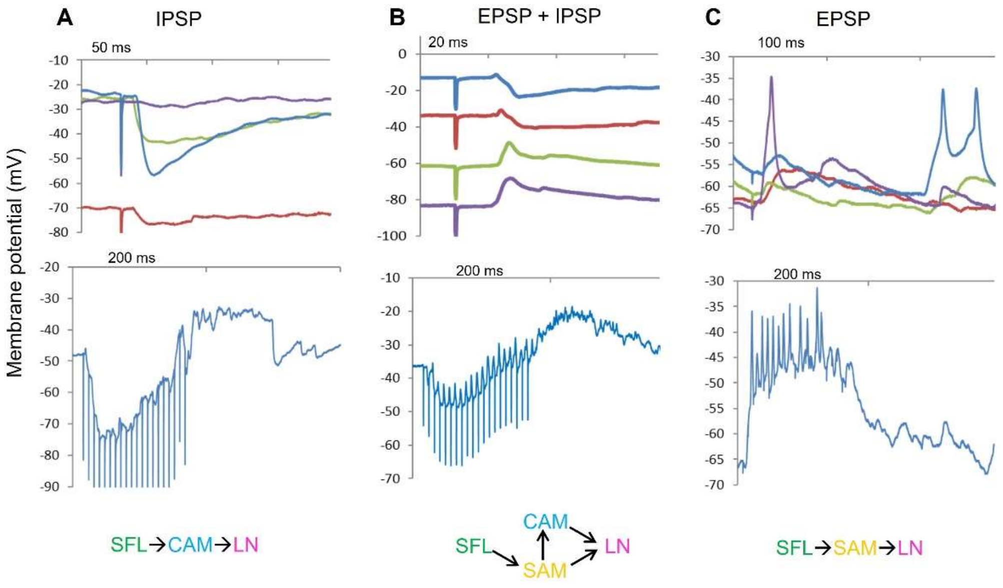


**Figure 7 - figure supplement 1.** Three examples for previous whole cell LN recording that support the interpretations of the three types of connectivity pathways revealed by the current study (shown at the bottom of the respective A,B and C panels). A. An intermitted IPSP is evoked by a single pulse stimulation of the SFL tact. The red trace shows the expected decline in the IPSP amplitude as a result of hyperpolarizing the membrane potential to around -70mV. The lower blue trace shows the summation of single IPSPs evoked by a train of 20 pulses at 50 Hz. The IPSPs are depressed as the train continues. B. An example for LN in which SFL stimulation evoked a profound EPSP followed by an IPSP. Changing the membrane potential by injecting constant current facilitates discrimination between the excitatory and inhibitory phases of the PSP. The lower blue trace shows the summed postsynaptic response evoked by a train as in A. C. An example that shows a LN that received an intermitted EPSP that occasionally reached the action potential threshold (note that the action potential doesn’t cross the zero membrane potential, indicating passive propagation of a remotely generated spikes back to the cell body, typical for invertebrates monopolar neurons). The lower blue trace panel shows robust excitatory summation of a train of EPSPs.

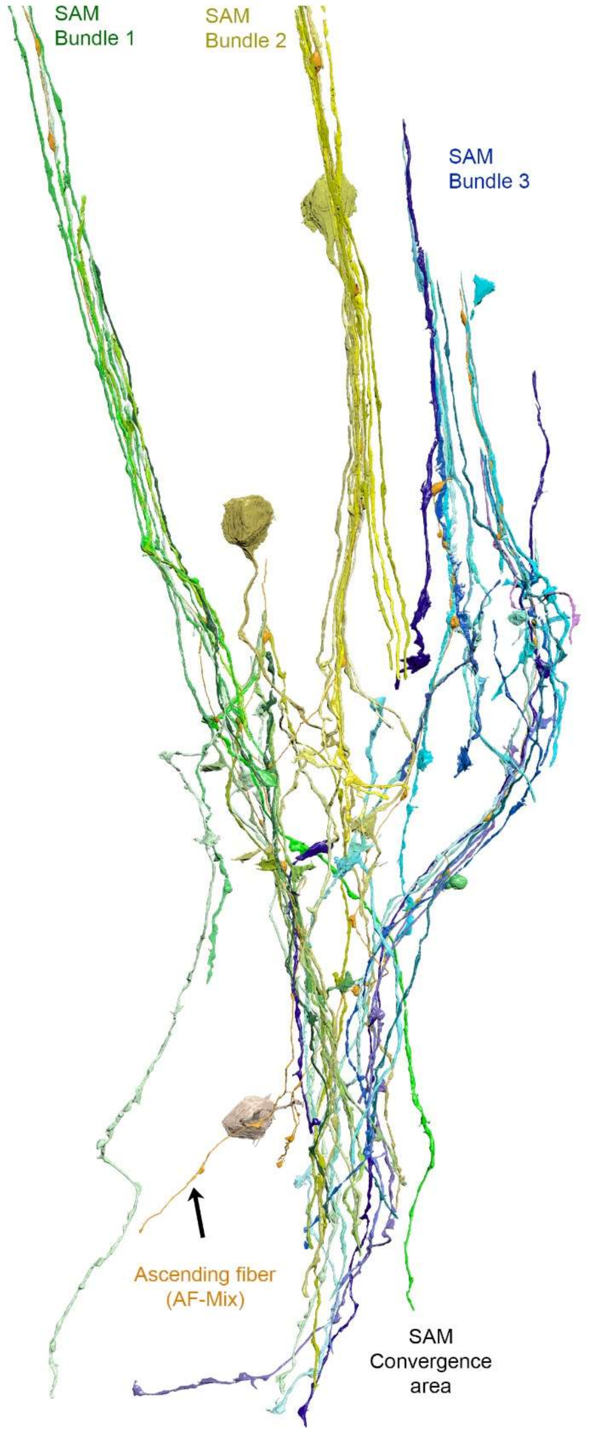


**Figure 9 - figure supplement 1**. **Reconstruction of one AF-mix (orange) and its inputs from the SAMS (yellow).** Notice that the AF intermingled with the SAMs. Within the cortex, the SAMS innervating the AF form distinct trunks/bundles that appear to converge within the neuropil. The SAMs innervating this AF appear to converge within the neuropil.
